# Single-Cell Transcriptomic Analyses of Tumor Ecosystems and Spatial Architectures in Human Small Cell Lung Cancer

**DOI:** 10.1101/2022.01.19.475968

**Authors:** Huiling Ouyang, Jian Chen, Chaoliang Xu, Cong He, Luoyan Sheng, Jing Wang, Deshen Pan, Jun Lin, Arnaud Augert, Yuming Zhu, Deshui Jia

## Abstract

Small cell lung cancer (SCLC) is a highly heterogenous disease characterized by aggressive phenotypes and poor prognosis. In order to dissect the cellular components and their spatial distribution in SCLC ecosystems, here we performed single-cell RNA sequencing and spatial transcriptomics analyses of 12 resected human primary SCLCs. Our analysis revealed extensive heterogeneity of both tumor and non-tumor cells, and a predominant immunosuppressive microenvironment. Importantly, multiple hybrid tumor cell states were unmasked, including hybrid SCLC and lung adenocarcinoma cells as well as hybrid tumor and immune cells, indicating high phenotypic plasticity of tumor cells. Moreover, we revealed the cellular origin and evolution of these hybrid tumor cells, and critical roles of tumor-stromal cellular crosstalk in driving the development of hybrid cells. In conclusion, this study deepens our understanding of human SCLC ecosystems and provides rationales to target the hybrid tumor cells and immunosuppressive microenvironment for SCLC therapies.

## INTRODUCTION

Small cell lung cancer (SCLC) has been a refractory type of lung cancer characterized by fast relapse after initial response to chemotherapy(1). Unfortunately, treatment choices have been very limited for SCLC patients in the last three decades(2). Even worse, lack of surgical specimens for biomedical research has been a barrier, and many studies heavily rely on preclinical models, such as genetically engineered mouse and patient-derived xenograft (PDX) models(3,4). However, tumors derived from these models cannot fully recapitulate the biological characteristics of human SCLCs such as infiltrating immune cells and stromal cells and spatial architectures of tumor ecosystems. Notably, SCLC has been revealed as a highly heterogenous disease, showing extensive inter- and intra-tumoral heterogeneity in both murine and human SCLC tumors, which directly associates with the aggressive phenotypes and poor prognosis(5-7). Recently, four molecular subtypes of SCLC have been defined based on the differential expression of transcription factors (ASCL1, NEUROD1, POU2F3 and YAP1), including SCLC-A and SCLC-N neuroendocrine (NE) subtypes, as well as SCLC-P and SCLC-Y non-NE subtypes(8). Different SCLC subtypes have distinct vulnerabilities to targeted therapies(9). For example, SCLC-A subtype was observed to be more sensitive to BCL2 inhibitors while the SCLC-N and SCLC-P subtypes were more sensitive to Aurora kinase inhibitor and PARP inhibitor, respectively(9). Recently, an inflamed subtype (SCLC-I) which exhibits increased sensitivity to immune checkpoint blockade (ICB) was identified(9,10). Additionally, MHC I expression which is increased in a subset of SCLC cells with non-NE feature, has been identified as a predictor of ICB efficacy(11). Given the heterogenous nature of SCLC tumors, there is an urgent need to dissect the tumor compositions and provide a more refined classification of SCLC subtypes in order to develop the next generation of precision therapies.

Recently, single-cell RNA sequencing (scRNA-seq) has been widely used to investigate tumor heterogeneity in human cancers, including non-small cell lung cancer, pancreatic cancer, head and neck squamous cell carcinoma and breast cancer, which has provided an unprecedent resolution to characterize the molecular features of individual cell and cell-cell communications among distinct cellular components in the tumor ecosystems(12-16). Remarkably, scRNA-seq has also been employed to investigate the tumor heterogeneity of SCLC using PDX, CDX models and human specimens(17-19). For example, a PLCG2-high tumor cell population with stem-like features was uncovered by scRNA-seq analyses of 21 human SCLC specimens(20). However, most of the SCLC specimens sequenced in these studies are biopsy samples and the number of individual cells sequenced per sample is relatively limited. Moreover, the landscape of infiltrated immune cells and stromal cells remains mostly unexplored. In addition, the spatial architectures of distinct cellular components in human SCLC ecosystems have not yet been investigated. To address these limitations, this study applied combined scRNA-seq and spatial transcriptomics analyses to dissect the cellular components and their spatial distribution in human SCLC tumors. Our findings revealed extensive tumor heterogeneity within tissues and tumors. Our work also revealed a predominant immunosuppressive microenvironment in human SCLC ecosystems. Importantly, multiple hybrid tumor cell states were uncovered, including a subset of hybrid tumor and immune cells. By integrating scRNA-seq and spatial transcriptomics analyses, we depicted the distribution of distinct clusters and their interactions in individual tumors. Altogether, our work provides a single-cell atlas and spatial distribution of the cellular components in SCLC ecosystems.

## RESULTS

### A Single-Cell Transcriptome Atlas of Human Primary SCLC Tumors

In order to dissect the heterogenous cellular components and their spatial distribution in SCLC ecosystems, a total of 12 primary tumors were collected from 12 SCLC patients who underwent surgery (Supplementary Table 1). First, we performed scRNA-seq analyses of 7 SCLCs, including 3 chemotherapy-treated and 4 untreated tumors (Figure 1A). In order to identify rare cell subpopulations in each tumor, we sequenced an average of ∼13,800 single cells per tumor resulting in a total of 81,841 high-quality cells across all tumors (Figure 1B, Supplementary Table 2). Through unsupervised clustering analysis, we identified 27 distinct cell clusters, including tumor cells, leukocytes, stromal fibroblasts and endothelial cells (Figure 1C). We further inferred copy number alterations (CNV) for each cluster in order to distinguish the tumor from non-tumor cells (Supplementary Figure 1A). Accordingly, based on the inferred CNV and expression of canonical cell type markers, 15 clusters were defined as tumor cells, whereas 9 and 3 clusters were classified as leukocytes and stromal cells, respectively (Figure 1C-D, Supplementary Figure 1B). Remarkably, we observed that a large number of tumor cells express both ASCL1 and NEUROD1, showing features of a hybrid state between SCLC-A and SCLC-N subtypes (Figure 1D). Furthermore, three major types of stromal cells were identified in the tumor microenvironment (TME), including FAP+ ACTA2+ myofibroblasts, MCAM+ ACTA2+ perivascular-like cells and PECAM1+ CD34+ endothelial cells (Figure 1D). Moreover, we observed extensive inter- and intra-tumoral heterogeneity across the 7 SCLCs (Figure 1E-F, Supplementary Figure 1C). Notably, we found that the numbers of infiltrated immune cells, stromal fibroblasts and endothelial cells are relatively low in the SCLC ecosystems (Figure 1G). In addition, we observed that the proportions of infiltrated immune cells and stromal cells are relatively low in chemotherapy-treated tumors compared with untreated tumors, indicating that chemotherapy may remodel the SCLC microenvironment (Supplementary Figure 1D-E).

**Figure 1.**
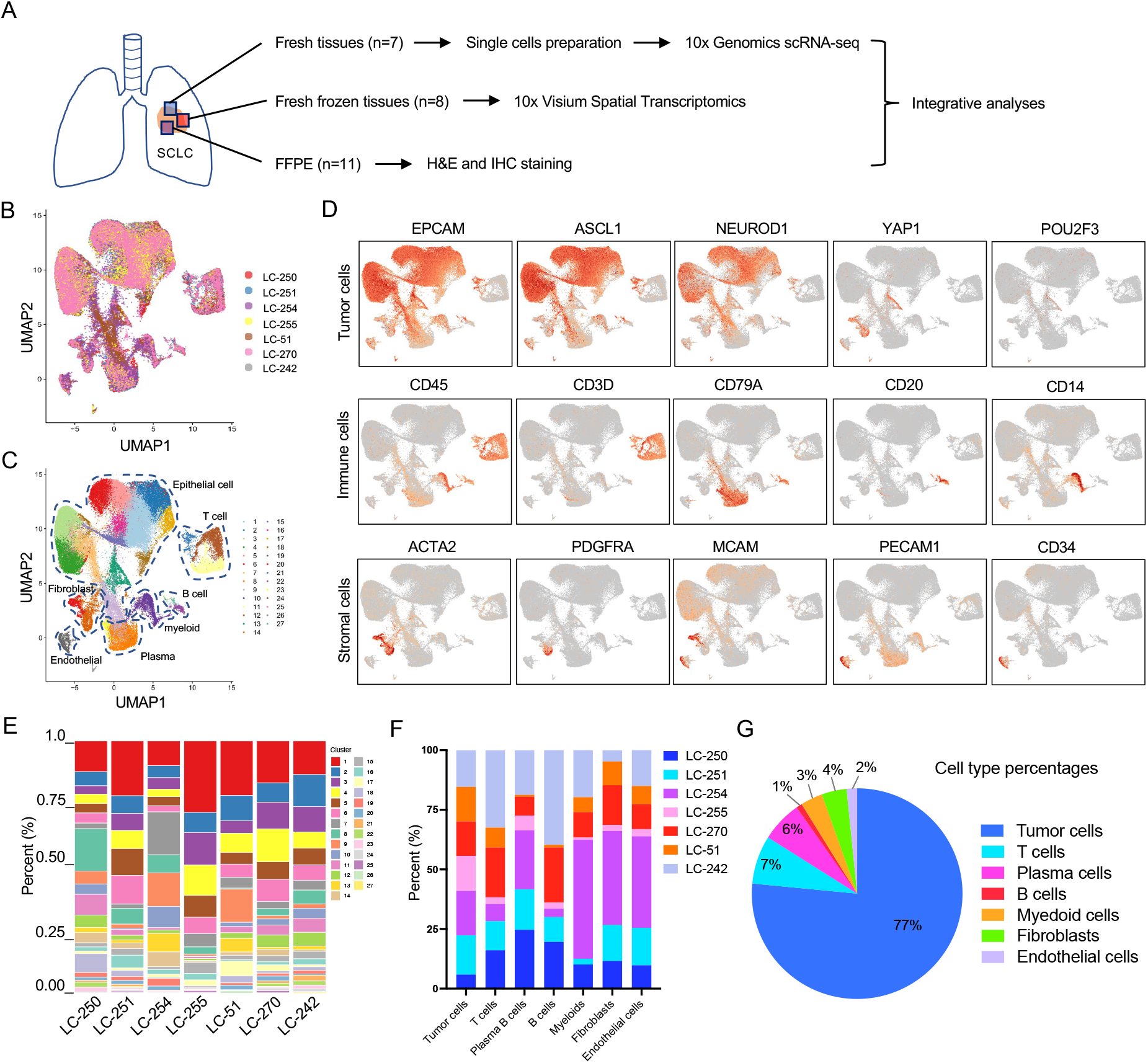
Single-cell atlas of cellular components in human SCLC ecosystems. A, Flowchart showing the study design and workflow. B, UMAP visualization of all 81,841 cells from scRNA-seq analyses colored by samples. C, UMAP visualization of all cells colored by clusters. Clusters were defined by graph-based clustering analysis. Cell types were annotated based on canonical markers. D, UMAP plots showing the relative expression of representative cell lineage markers in all cell clusters. Color bar refers to the log2 expression levels of each gene. E-F, The proportions of each cell cluster (F) and cell type (G) in individual SCLC sample. G, Percentage of each cell type in 7 SCLC ecosystems.

### Spatial Transcriptomic Landscape of Cellular Components in SCLC Ecosystems

In order to explore the spatial localization of distinct cellular components and their interactions, we next performed spatial transcriptomics analyses on 8 immunohistochemically validated human SCLC tumors, including 6 SCLC-A, 1 SCLC-N and 2 SCLC-P tumors (Figure 2A, Supplementary Figure 2A). Notably, 3 (LC-250, LC-251 and LC-51) of the 8 tumors were sequenced by both scRNA-seq and spatial transcriptomics (Supplementary Table 1). A total of 16,853 spots from 8 tumors were captured, sequenced and grouped into 17 clusters (Figure 2B, Supplementary Table 2). Next, we explored the spatial distribution of individual subpopulations within each tumor (Figure 2C, Supplementary Figure 2B). Consistently, the extensive inter- and intra-tumoral heterogeneity of SCLCs was further revealed (Supplementary Figure 2C). Therefore, we subsequently analyzed the spatial transcriptomics data of individual tumor separately.

**Figure 2.**
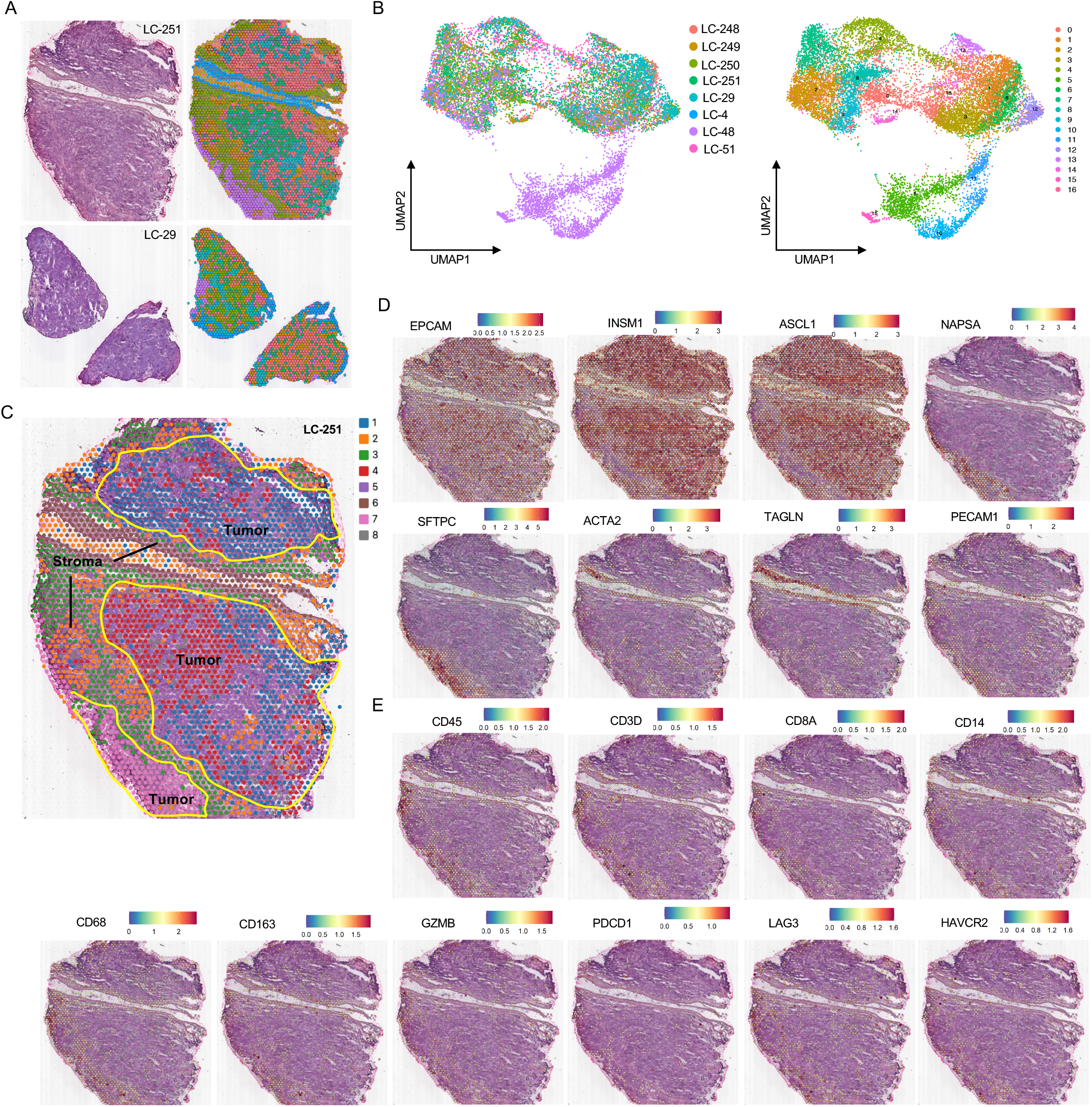
Spatial transcriptomic analyses revealed physical distribution patterns of cellular components in SCLC ecosystems. A, Images of H&E staining and spatial spot clusters for LC-251 and LC-29 tissue regions. B, UMAP visualization of 16,853 spots colored by samples and clusters, respectively. C, Spatial feature plots of 8 clusters and their distribution within the LC-251 tissue section. D-E, Spatial feature plots showing the expression of representative markers (E), immune cells and exhausted markers (E) in all cell clusters within the LC-251 tissue. Color bar refers to the relative expression levels of each gene.

For instance, 8 spatial spot clusters were identified in the LC-251, including 5 tumor cell clusters (clusters 1, 4, 5, 7 and 8) and 3 stromal cell clusters (clusters 2, 3 and 6). We observed that clusters 1, 4 and 5 are located in the tissue center whereas cluster 7 is located at the tissue edge within LC-251 tissue (Figure 2C). By interrogating differential gene expression of each cluster, we found that cluster 4 show high expression of H3F3A, HMGN2 and TXN, whereas cluster 7 exhibit high expression of SFTPC and NAPSA (Figure 2D, Supplementary Figure 2D). Importantly, we observed that cluster 7 expresses markers of both SCLC and lung adenocarcinoma (LUAD), showing features of hybrid transcriptional state (Figure 2D). In addition, we found that cluster 5 exhibit high expression of stromal cell markers such as COL4A1, SPARC, ESM1 and RGS5 (Supplementary Figure 2D). Similarly, hybrid SCLC/LUAD cells were also identified in the LC-51 tissue (Supplementary Figure 2E).

Next, we examined the spatial distributions of stromal and immune cells within SCLC tissues. Within the LC-251 tissue, we observed that stromal cells (clusters 2 and 3) surround and separate the different tumor cell clusters (Figure 2C). Moreover, CD45+ immune cells tend to distribute sparsely within the tissue, and are mostly exhausted CD8+ T cells and M2 macrophages (Figure 2E), indicating an immunosuppressive microenvironment. To our knowledge, this is the first analysis that explores the spatial architectures of cellular components in human SCLC ecosystems using high-resolution transcriptomics. Our results not only reveal the extent of intra-tumoral heterogeneity but also provides a spatial distribution of distinct cell types and their molecular features within SCLC tumors.

### Immune Cell Landscape and Spatial Distribution in SCLC Ecosystems

The diversity and transcriptomic features of infiltrated leukocytes and stromal cells in human SCLC ecosystems are relatively unexplored. In this study, a total of 14,755 infiltrating leukocytes were identified by scRNA-seq. We observed high inter- and intra-tumor heterogeneity of immune cells across the 7 SCLCs (Supplementary Figure 3A-B). For example, LC-242 and LC-254 tumors have relatively high infiltration of immune cells while LC-51 and LC-255 tumors have low infiltration. We further re-clustered all immune cells into 23 subpopulations (Figure 3A). Based on the expression of canonical markers, most subpopulations could be defined as CD4 T cells, CD8 T cells, NKT cells, plasma and B cells, Tryptase+ mast cells, plasmacytoid dendritic cells (pDCs) and monocytes/macrophages (Figure 3B-C). However, 5 subpopulations (clusters 7, 11, 12, 13 and 22) were observed to also express markers of epithelial and neuroendocrine cells, such as EPCAM, INSM1 and ASCL1, showing features of hybrid tumor and immune cells (Figure 3B-C). We postulated that these cells might be tumor cells expressing immune cell markers, which is further investigated in the following section. Of note, positive expression of immune cell markers on tumor cells has been described in several cancers(21-23).

**Figure 3.**
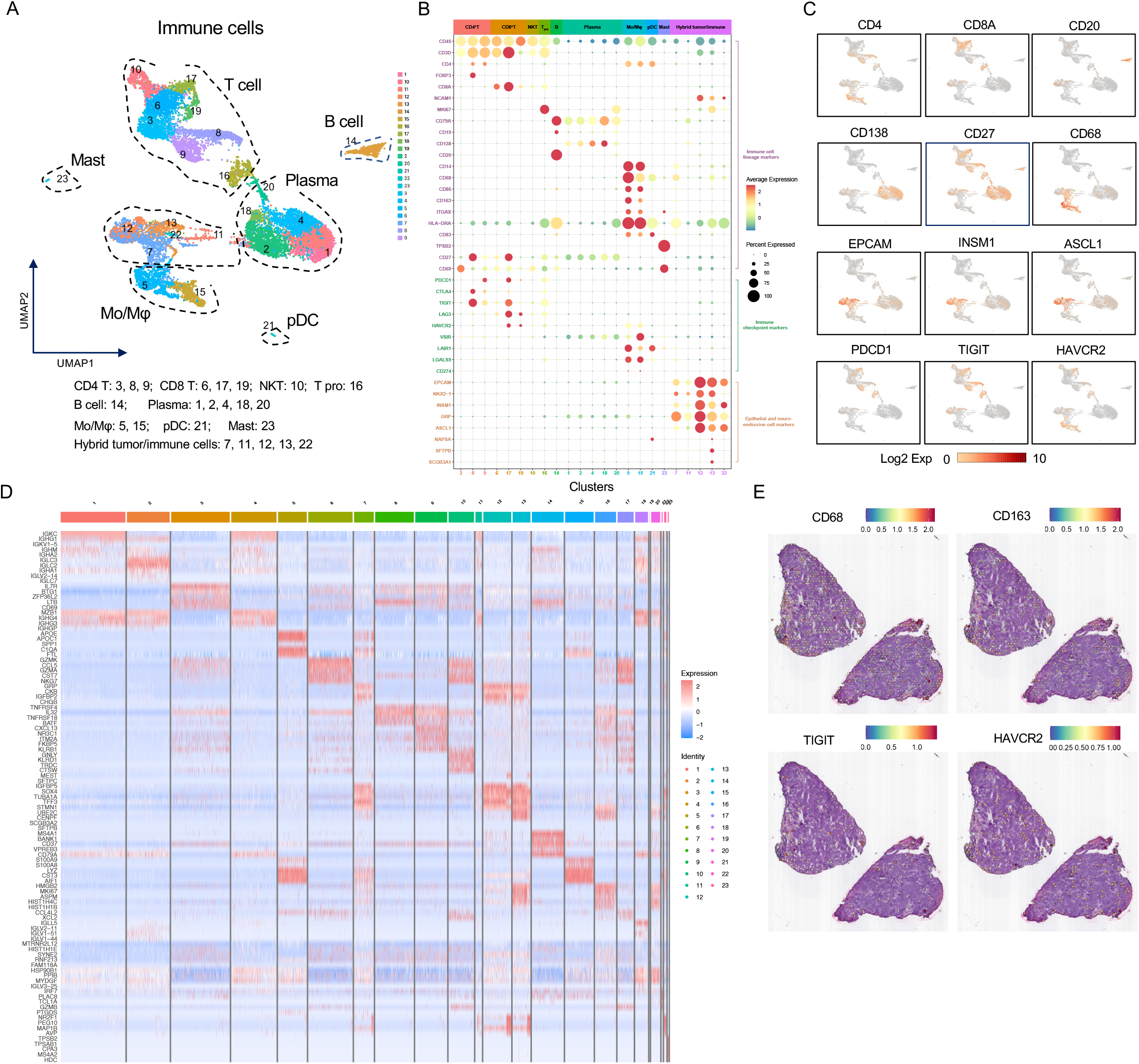
Single-cell landscape and spatial distribution of immune cells in human SCLC ecosystems. A, UMAP visualization of all immune cells colored by clusters. Cell types were annotated based on canonical immune cell markers. B, Dot plot showing the expression of representative markers for each immune cell cluster grouped by cell types. Color bar refers to the expression values, dot size means the proportion of positive gene expression in individual cluster. C, UMAP plot showing the expression of representative markers for immune cells. Color bar refers to the log2 expression levels of each gene. D, Heatmap showing the top 5 upregulated genes in each of the 23 immune cell clusters. E, Spatial feature plots showing the expression of macrophages and exhausted markers within the LC-29 tissue. Color bar refers to the expression levels of each gene.

Next, we characterized the T cell subpopulations within the SCLC ecosystems. For CD4+ T cells, cluster 8 represented regulatory T cells characterized by high expression of FOXP3, cluster 9 represented follicular helper T cells showing high levels of CXCL13, NR3C1 and FKBP5, whereas cluster 3 referred to cytotoxic T cells exhibiting high levels of IL7R and BTG1 (Figure 3D). Regarding CD8+ T cells, clusters 6, 17 and 19 represented cytotoxic T cells with high expression of GZMK, CCL5, NKG7 and GZMA, whereas cluster 10 expressed high levels of GNLY and KLRD1, which represents NKT cells (Figure 3D). In addition, we found that cluster 16 contains both CD4+ and CD8+ T cells and exhibits high expression of MKI67 and ASPM genes, representing a subpopulation of proliferating T cells (Figure 3D). Importantly, we found that most T cells exhibit positive expression of immune checkpoint genes associated with T cell exhaustion, such as PDCD1, CTLA4, TIGIT, LAG3 and HAVCR2 (Figure 3B-C), indicating that an immunosuppressive microenvironment is predominant in the SCLC microenvironment.

Previously, T cells and macrophages have been observed to be the most abundant infiltrated immune cells in SCLC, whereas B cells were sparse as assessed by immunohistochemical analysis(24). Here, we found that the proportion of infiltrated B cells is comparable to T cells and myeloid cells in the SCLC ecosystems, but the majority of B cells are plasma cells and exhibit remarkable heterogeneity (Figure 3A-B). We showed that plasma cells are characterized by high expression of IGKC, IGHG1 and MZB1 whereas CD20+ B cells are characterized by high expression of BANK1 and CD37 (Figure 3D). Moreover, we found that the CD20+ B cell population is composed of both activated (CD19+ CD20+ CD69+ CD27+) and exhausted (CD19+ CD20+ CD69+ CD27-) cells (Figure 3B-C). Through functional annotation of differentially expressed genes, we observed that antigen processing and presentation is among the top enriched pathways in CD20+ B cells (Supplementary Figure 3C). Moreover, we observed that both plasma and CD20+ B cells distribute sparsely within tissues (Supplementary Figure 3D).

In addition, we identified 4 subpopulations of myeloid cells. The cluster 5 represented C1QB+ macrophages characterized by high expression of APOE, APOC1 and SPP1, whereas cluster 15 represented FCN1+ monocytes showing high expression of S100A9, LYZ and CST3 (Figure 3B-D). Besides, we observed that cluster 23 represents a subpopulation of mast cells with high expression of TPSB2 and TPSAB1, whereas cluster 21 represents pDCs with high expression of IRF7 and GZMB (Figure 3B-D). Moreover, we observed that most macrophages exhibit high expression of CD163, LAIR1, HAVCR2, LGALS9, VSIR and LAG3, indicating that immunosuppressive M2 macrophages are predominant in SCLC TME (Figure 3B-C). Finally, we observed that macrophages are predominant and distribute primarily within the stroma of LC-251 and LC-29 tumors (Figure 2E, Figure 3E). Altogether, our analyses provide a single-cell atlas of infiltrating immune cells and their spatial distributions in human SCLC microenvironment.

### Landscape of Stromal Fibroblasts and Endothelial Cells in SCLC Ecosystems

Cancer-associated fibroblasts (CAFs) are major components of the TME and have been shown to play critical roles in tumor initiation and progression(25). However, little is known about the categories of CAFs and their functional roles in SCLC. Here, a total of 3,145 CAFs were identified by scRNA-seq, which were grouped into 15 subpopulations (Figure 4A-B). Furthermore, 5 states of fibroblasts were revealed in SCLC ecosystems (Figure 4C). S1 CAFs resembled perivascular-like cells, showing high expression of MCAM and ACTA2, enriched for focal adhesion, cGMP-PKG signaling and vascular smooth muscle contraction pathways (Figure 4D-E, Supplementary Figure 4A). S2 CAFs represented myofibroblasts characterized by high expression of FAP and PDGFRA, enriched for ECM-receptor interaction and focal adhesion pathways (Figure 4D-E, Supplementary Figure 4B). Importantly, S3, S4 and S5 fibroblasts resembled tumor cells undergoing an EMT program, co-expressing markers of both mesenchymal and neuroendocrine cells, including ACTA2, INSM1, ASCL1 and GRP (Figure 4D-E, Supplementary Figure 4C-D). Accordingly, we could classify fibroblasts into three major subtypes, including FAP+ myofibroblasts, MCAM+ perivascular-like cells, and EPCAM+ fibroblasts. Antigen-presenting and inflammatory CAFs have been identified in multiple cancers(26). Interestingly, we found that a subset of EPCAM+ CAFs also express markers of immune cells, such as high expression of CD24, IGHG2 and IGLC2 in cluster 3, as well as high expression of IGHG1, IGHG4 and MZB1 in cluster 13 (Supplementary Figure 4E). Collectively, our study provides a single-cell atlas of CAFs in human SCLC ecosystems and highlights that the EPCAM+ CAFs most likely derive from tumor cells that have undergone an EMT program.

**Figure 4.**
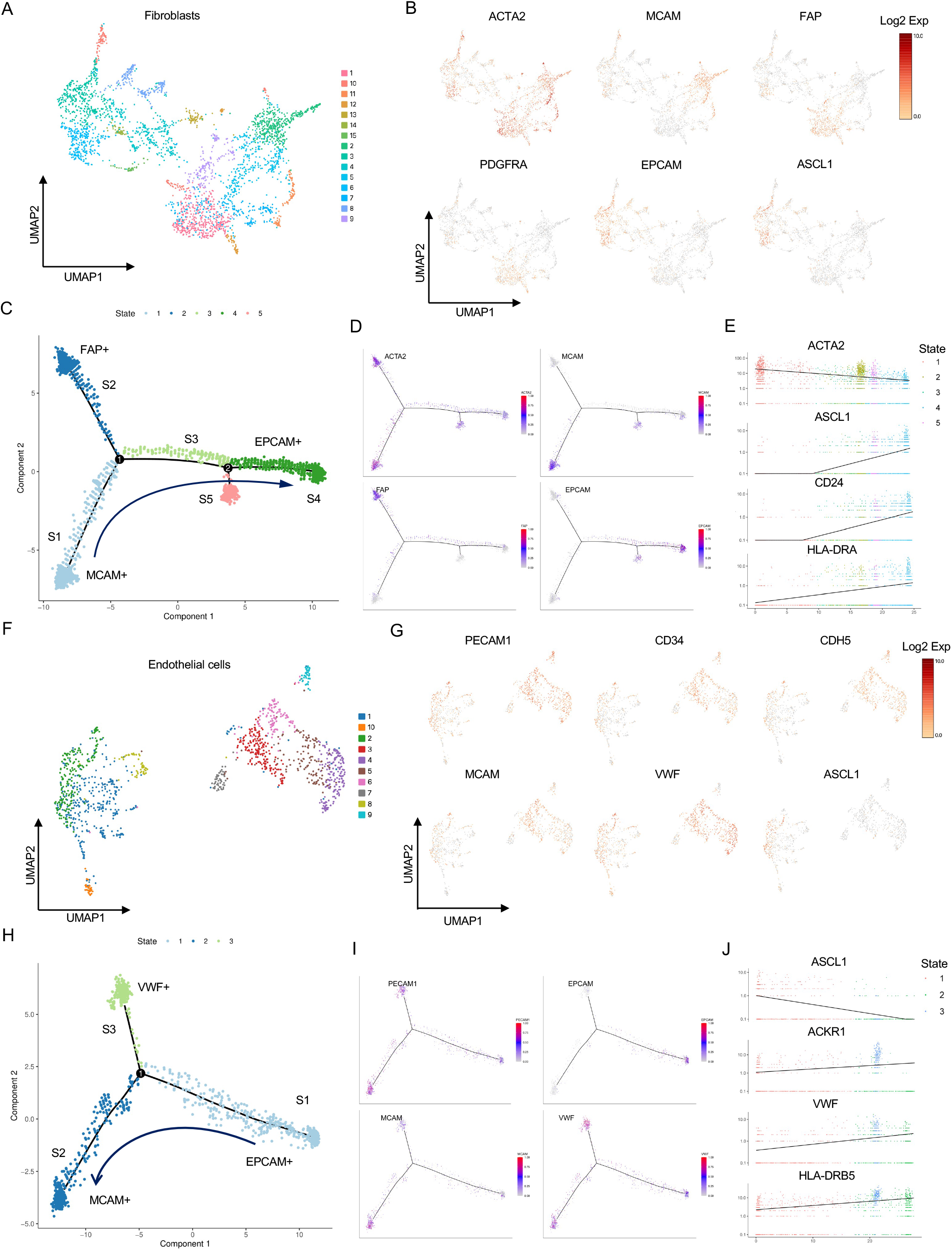
Single-cell landscape and spatial distribution of stromal cells in human SCLC ecosystems. A, UMAP visualization of all fibroblasts colored by clusters. B, UMAP plots of representative markers for fibroblasts (ACTA2, MCAM, FAP and PDGFRA) and epithelial/neuroendocrine cells (EPCAM and ASCL1). Color bar refers to the log2 expression levels of each gene. C, States of fibroblasts in human SCLC ecosystems. D, The expression of representative genes in each state of fibroblasts. E, Dot plot showing the expression of representative genes in each state of fibroblasts. F, UMAP visualization of all endothelial cells colored by clusters. G, UMAP plots showing representative markers of ECs (PECAM1, CD34, CDH5, MCAM and VWF) and neuroendocrine cells (ASCL1). Color bar refers to the log2 expression levels of each gene. H, States of ECs in human SCLC ecosystems. I, The expression of representative genes in each state of ECs. J, Dot plot showing the expression of representative genes in each state of ECs.

In addition, we identified a total of 1,308 vascular endothelial cells (ECs) in SCLC ecosystems, which contained 10 subpopulations and could be grouped into 3 states (Figure 4F-H). S1 ECs resembled tumor-endothelial transition cells, co-expressing both endothelial and neuroendocrine cell markers, such as PECAM1, ASCL1 and GRP (Figure 4I-J, Supplementary Figure F). S2 ECs showed typical features of endothelial cells, expressing high levels of MCAM, ANGPT2 and ESM1, whereas S3 ECs exhibited high expression of VWF, ACKR1, HLA-DRB5 and RGS5 (Figure 4I-J, Supplementary Figure 4G). Notably, it has been reported that non-neuroendocrine SCLC cells are transcriptionally primed to undergo vascular mimicry and express gene expression profiling of pseudohypoxia(27). Collectively, these findings suggest that one of the mechanisms by which SCLC cells induce angiogenesis to support their rapid growth may be through tumor-endothelial transition. Finally, we highlight the heterogeneity of stromal cells and their spatial localization in human SCLC tissues (Supplementary Figure 4H).

### Hybrid Tumor Cells with Multiple Cell Lineage Features Revealed in Human SCLCs

In this study, a total of 62,633 tumor cells were identified by scRNA-seq accounting for ∼77% of all cellular components in the 7 SCLC ecosystems (Figure 1G). We further re-clustered these tumor cells into 26 subpopulations, which exhibit extensive heterogeneity within tissues and tumors (Figure 5A, Supplementary Figure 5A). We observed that most subpopulations express both ASCL1 and NEUROD1, showing feature of hybrid SCLC-A/N subtypes (Figure 5B-C). We also observed a subpopulation of YAP1+ cells (cluster 21) displaying high expression of LUAD markers such as NAPSA, SFTPB and SCGB3A1 (Figure 5B). Moreover, we observed a few POU2F3+ tumor cells, which sparsely distribute in multiple clusters (Figure 5B, Supplementary Figure 5B). Remarkably, in addition to the hybrid SCLC-A/N cells, we identified multiple other hybrid tumor cell states, including hybrid Epithelial/Mesenchymal (E/M) cells, SCLC/LUAD cells and tumor/immune cells. For instance, we observed that cells in clusters 21, 16, 11, 17, 25 and 8 show hybrid E/M features, co-expressing both epithelial and mesenchymal markers (Figure 5B-C). Interestingly, a subset of cells with EMT features also expressed LUAD markers, especially in the LC-51 sample (Figure 5B-D), indicating that hybrid SCLC/LUAD cells may undergo an EMT program. Moreover, we found that some tumor cells exhibit high expression of immune cell markers (CD45, CD27 and HLA-DRA), showing hybrid tumor/immune features (Figure 5B-C). Importantly, we confirmed the presence of tumor cells expressing immune cell markers in several human SCLC cell lines by flow cytometry analyses (Figure 5E, Supplementary Figure 5C).

**Figure 5.**
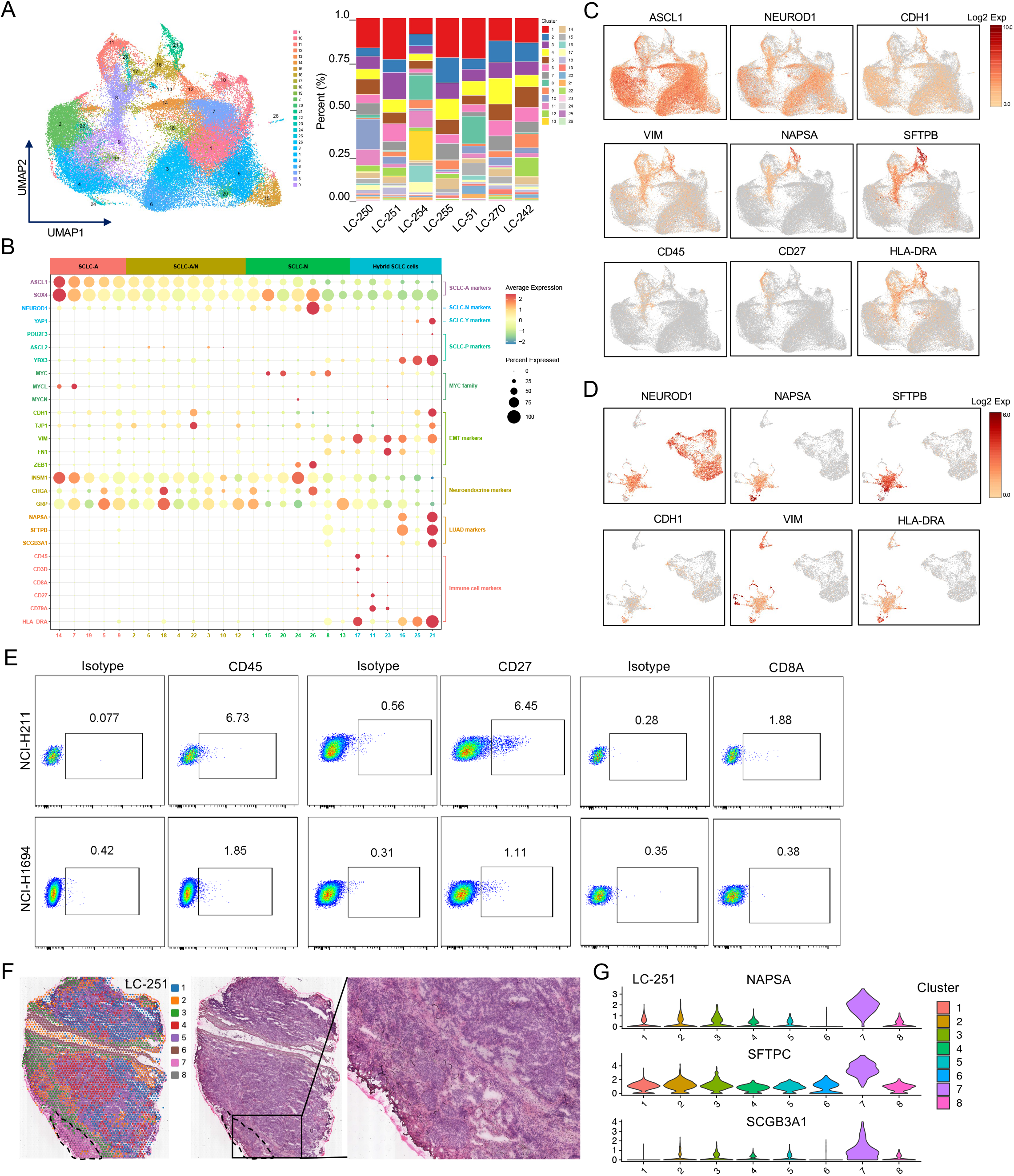
Single-cell transcriptome analysis revealed multiple hybrid states of SCLC cells. A, UMAP visualization of all 62,633 tumor cells colored by clusters. The relative proportion of individual clusters in each sample was illustrated. B, Dot plot showing the expression of representative markers for each immune cell cluster grouped by cell types. Color bar refers to the expression values, dot size means the proportion of positive gene expression in individual cluster. C, UMAP plots of representative markers in all tumor cell clusters across 7 samples. Color bar refers to the log2 expression levels of each gene. D, UMAP plots of representative markers in all cell clusters of the LC-51 tumor. Color bar refers to the log2 expression levels of each gene. E, Flow cytometry results showing the proportion of tumor cells expressing each of the immune cell markers (CD45, CD27 and CD8) in human NCI-H211 and NCI-H1694 cell lines, respectively. F, Images of spatial spot clusters and H&E staining for the LC-251 tissue. G, Violin plot showing the relative expression of NAPSA, SFTPC and SCGB3A1 in each spatial spot cluster of the LC-251 tumor.

Recently, high MHC I expression was observed in a subset of SCLCs with non-neuroendocrine features and associated with durable response to immune checkpoint blockades (ICB), suggesting that MHC I could be used as a predictive biomarker of immune response for SCLC patients(11). Here we observed positive expression of genes encoding MHC I molecules (HLA-A, HLA-B and HLA-C) in almost all SCLC tumor cells (Supplementary Figure 5B). Interestingly, we found that the expression of genes encoding MHC II molecules (HLA-DRA and HLA-DRB1) was also upregulated in clusters 21, 16, 17, 11, 8 and 12 (Figure 5B-C, Supplementary Figure 5B). The MHC II molecules are typically expressed on professional antigen-presenting cells, such as DCs, monocytes and B cells. Collectively, these findings indicate that most hybrid tumor/immune cells also exhibit EMT and/or hybrid SCLC/LUAD cell features and may be more sensitive to ICB therapy. Similarly, we also identified a subset of tumor cells expressing endothelial cell markers, such as PECAM1, CDH5 and CD34 (Supplementary Figure 5D).

Finally, our spatial transcriptomics analyses revealed that hybrid SCLC/LUAD cells mainly locate at the tumor edge within the LC-251 tissue whereas they are located within the center of the LC-51 tissue (Figure 5F-G, Supplementary Figure 2E). The hybrid SCLC-A/SCLC-P cells in the LC-4 tumor and hybrid SCLC-P/SCLC-Y cells in the LC-29 tumor were randomly distributed within tissues (Supplementary Figure 5E-F). Taken together, our work reveals multiple hybrid tumor cells in human SCLCs.

### Trajectory Analysis Reveals Cell of Origin of Hybrid SCLC Cells

In order to trace the evolution of tumor cells in SCLC ecosystems, we performed trajectory analysis of all tumor cells. We identified 6 branches and 9 cell states (Figure 6A). We observed that branch 1 is composed of states 1, 2, 3 and 4 cells, whereas branches 2, 3, 4, 5 and 6 are composed of states 9, 8, 7, 5 and 6 cells, respectively. Notably, cells in the branches 5 and 6 were mainly derived from the LC-255 tumor, suggesting a more differentiated state (Figure 6A). We observed that the evolutionary trajectory starts from the state 1 cells. As time progresses, cells in the other 8 states evolved along the trajectory. We observed 4 branch points during the evolution of SCLC cells, and all of them could successfully differentiate into distinct branches (Figure 6A). Furthermore, we analyzed the expression of canonical markers of SCLC and LUAD that change over pseudotime (Figure 6B). We observed the lowest expression of ASCL1, NEUROD1 and INSM1 at the beginning of the trajectory (state 1), whereas the expression of these genes increased stepwise over pseudotime (Figure 6B). In contrast, we found that the expression of LUAD and mesenchymal markers (NAPSA, SFTPC, VIM) decrease stepwise over pseudotime (Figure 6B). Consistently, we observed that state 1 cells show features of hybrid SCLC/LUAD cells, whereas state 9 cells represent hybrid SCLC-A/SCLC-N cells, pointing to a potential cell lineage transition between LUAD and SCLC (Figure 6B). Together, our findings suggest that one potential cell of origin of hybrid tumor cells, such as hybrid SCLC/LUAD cells, might originate from a LUAD cell lineage.

**Figure 6.**
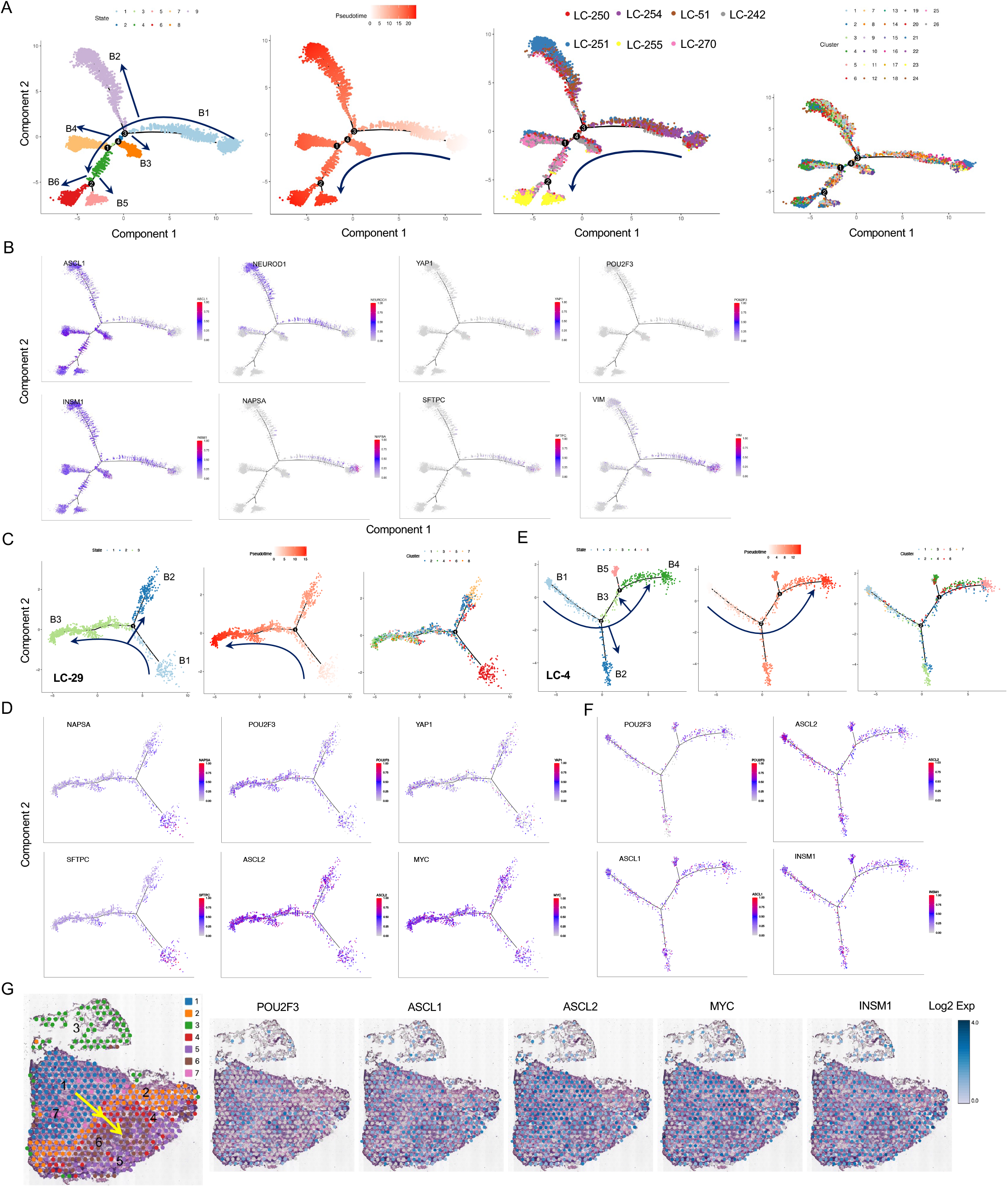
Trajectory analysis revealed dynamic evolution and cell origins of hybrid SCLC cells. A, Trajectory analysis of tumor cells. The cells are colored by differentiation states, pseudotime, samples and cell clusters, respectively. B, Expression of representative markers projected onto pseudotime space. C, Trajectory analysis of all cells in the LC-29 tumor using Spatial Transcriptomics data. The cells are colored by differentiation states, pseudotime and cell clusters, respectively. D, Expression of representative markers projected onto pseudotime space in the LC-29. E, Trajectory analysis of all cells in the LC-4 tumor using Spatial Transcriptomics data. The cells are colored by differentiation states, pseudotime and cell clusters, respectively. F, Expression of representative markers projected onto pseudotime space in LC-4. G, Spatial feature spots showing the evolution of distinct cell clusters and the expression of representative markers in the LC-4. Arrow shows the direction of evolution.

SCLC-P subtype accounts for ∼15% of all SCLC, and is recognized as a non-neuroendocrine type with more aggressive properties(8). It has been reported that some SCLC-P tumors could originate from rare tuft cells in lung epithelium and express canonical markers of tuft cells, such as POU2F3, ASCL2 and AVIL(28). However, the cellular origins of this subtype have not been well defined. In this study, we identified multiple hybrid SCLC-P cells, such as hybrid SCLC-A/SCLC-P cells in the LC-4 tumor and hybrid SCLC-P/SCLC-Y cells in the LC-29 tumor (Supplementary Figure 2F). In order to uncover new cell of origin for the SCLC-P subtype, we investigated the evolution of tumor cells in these tumors. Trajectory analysis revealed 3 branches and 3 cell states in the LC-29 tumor (Figure 6C). The trajectory root starts from the state 1 cells, which is composed by cells from cluster 6 and characterized by high expression of LUAD markers, such as the NAPSA, SFTPC (Figure 6D). As time progressed, tumor cells differentiated into 2 branches, whereby branch 2 was enriched with cells from cluster 2 and 7 while branch 3 was enriched with cells from cluster 1, 2, 3, 4, 5 and 8 (Figure 6C). Remarkably, we observed that the expression of LUAD markers decrease while the expression of POU2F3, YAP1, ASCL2 and MYC increase during the transition from branch 1 to branch 3 (Figure 6D). Moreover, we found that the LC-29 tumor expresses relatively high levels of KRT5, KRT8 and NKX2-1 genes (Supplementary Figure 6A). Together, these findings suggest that the SCLC-P subtype could originate from multiple cell types in the lung epithelium, such as keratin+ basal cells, alveolar cells and transformation from LUAD cells. In line with this hypothesis, a tuft cell-independent origin of SCLC-P has recently been revealed in the study of transformed SCLC(29).

In addition, we identified 4 branches and 5 cell states in the LC-4 tumor (Figure 6E). Based on the histological feature and gene expression signature of each cluster, we defined cluster 3 as adjacent normal lung cells, clusters 2 and 5 as stromal cells, and clusters 1, 4, 6 and 7 as tumor cells (Supplementary Figure 2). Accordingly, we observed that the trajectory root originates from state 1 cells, which was populated with tumor cells from clusters 1 and 7 and characterized by high expression of non-neuroendocrine markers, including ASCL2 and POU2F3 (Figure 6F). As time progressed, tumor cells could differentiate into states 4 and 5, which are composed by cells from clusters 4 and 6 and show high expression of neuroendocrine markers (Figure 6F). Therefore, we could visualize a transition from the SCLC-P to SCLC-A subtype within the LC-4 tumor, which starts from cluster 1 and ends at clusters 4 and 6 (Figure 6G). Together, these findings indicate that SCLC-P cells may differentiate into hybrid SCLC-A/SCLC-P cells.

### Exploration of Cellular Crosstalk in SCLC Ecosystems

Finally, in order to identify molecular mechanisms that drive tumor heterogeneity, especially the development of hybrid tumor cells, we analyzed ligand-receptor interactions among all the 27 cellular components in the 7 SCLC ecosystems. Accordingly, a total of 34,546 pairs of ligand-receptor interactions were identified (Figure 7A, Supplementary Table 3). Remarkably, we observed large number of interactions between tumor and immune cells. Myeloid cells exhibited the largest number of interaction pairs (Figure 7A). Most of interactions between myeloid and tumor cells are mediated by molecules associated with immune response and extracellular matrix, such as the CD74-MIF/COPA/APP, HLA-DPB1-TNFSF13B/NRG1 and SPP1-CD44 pairs, as well as mediated by chemokines and their receptors, such as the CCL3-IDE, CCL2-CCR10 and CXCL8-ACKR1 pairs (Figure 7B). Furthermore, we observed several known inhibitory interactions between T cells and tumor cells, such as the TIGIT-NECTIN2/NECTIN3 and PDCD1-CD274/PDCD1LG2 pairs (Figure 7C, Supplementary Table 3). It has been reported that CAFs can secrete numerous chemokines and growth factors, by which they were able to remodel the TME and regulate tumor progression(25). Here we observed extensive interactions between CAFs and tumor cells, most of which are mediated by chemokines and immune-related molecules, such as the TIMP1-FGFR2 and HLA-C-FAM3C pairs (Figure 7D). Similarly, we revealed interactions between endothelial and tumor cells, including the CD74-MIF and NRP1-VEGFB pairs (Supplementary Figure 6B-C).

**Figure 7.**
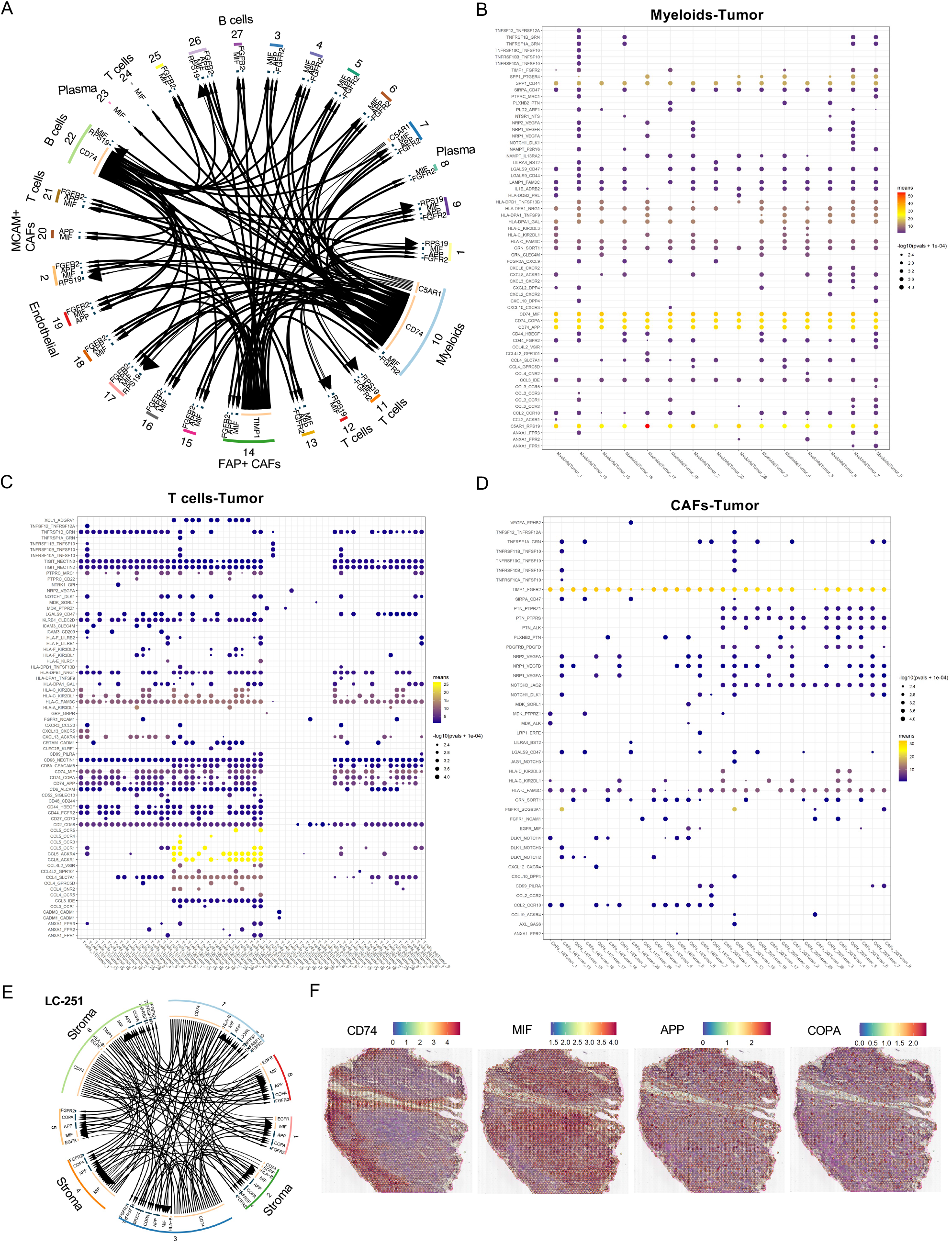
Identification of cell-cell communications within and across cellular components in human SCLC ecosystems. A, Circle plot illustrating the overview of ligand-receptor interactions among all 27 clusters of cellular components in human SCLC ecosystems. Clusters distinguished by colors, arrow lines start from ligands and end at receptors. B-D, dot plot showing the top interactions between myeloids and tumor cells (B), between T cells and tumor cells (C), and between CAFs and tumor cells (D). Dot colors represent the total means of individual partner average expression values, dot sizes represent the -log10(p values + 1e-04). E, Circle plot showing the interactions among 8 spatial spot clusters in the LC-251 tumor. Clusters distinguished by colors, arrow lines start from ligands and end at receptors. F, Spatial feature plots showing the expression of CD74, MIF, APP and COPA in all spatial spots of the LC-251.

Importantly, we identified multiple interactions between hybrid tumor cells and stromal cells. For example, we found that interactions between hybrid SCLC/LUAD cells (cluster 7) and myeloid cells are mediated by the CD74-MIF/COPA, C5AR1-RPS19 and SPP1-CD44 pairs (Figure 7B), whereas interactions between hybrid tumor/immune cells (cluster 9) and FAP+ CAFs (cluster 14) are mediated by chemokines, Notch ligands and their respective receptors (Figure 7D), indicating that stromal cells may drive the development of these hybrid tumor cells by enhancing cell-cell communication. Finally, we also displayed the cell-cell communication between different cell types within tissues. For example, we explored the interactions between stromal cells (clusters 2, 3 and 6) and tumor cells, especially hybrid SCLC/LUAD tumor cells (cluster 7) within the LC-251 tumor (Figure 7E, Supplementary Table 4). We observed that the expression of CD74 is relatively high in stromal cells, whereas expression levels of COPA, APP and MIF are relatively high in tumor cells (Figure 7F). Taken together, our work represents an initial step to dissect the intercellular communications between cellular components in the SCLC ecosystem, and provide a network of cellular crosstalk mediated by ligand-receptor interactions during SCLC progression.

## DISCUSSION

Understanding the cellular and molecular mechanisms that drive tumor heterogeneity and progression is critical to develop new therapeutic strategies. Single-cell RNA-seq has been used to analyze the tumor heterogeneity of SCLC(17,18,20,30). However, these studies have not provided a comprehensive single-cell atlas of all cellular components and their spatial architectures in human primary SCLC microenvironment. In particular the landscape of immune and stromal cells has not been thoroughly investigated. Here, we took an initial step to dissect the complex ecosystems of human SCLCs by using scRNA-seq and spatial transcriptomics technologies. SCLC is characterized as an immune cold tumor with low infiltration of immune cells(24). In accordance, our study also revealed low infiltration of immune cells in SCLC ecosystems. However, our data revealed the diversity and heterogeneity of infiltrated immune cells within tumors and tissues as well as their transcriptional signatures.

Importantly, our work demonstrated that an immunosuppressive microenvironment mediated by exhausted T cells and M2 macrophages is predominant in SCLC ecosystems. In addition, we explore the spatial distributions of distinct immune subpopulations within SCLC tissues and show that most immune cells sparsely localize within the SCLC tissues.

Furthermore, our study revealed an extensive heterogeneity of stromal fibroblasts in human SCLC ecosystems, as 15 subpopulations of CAFs were identified. Notably, we identified a subset of EPCAM+ fibroblasts, which may be tumor cells that underwent an EMT program. Recently, CD105 has been used as a marker to define stromal fibroblasts, and CD105− fibroblasts are able to promote anti-tumor immunity in pancreatic cancer(31). In the current study, we found that MCAM could distinguish fibroblasts in SCLC. The MCAM+ fibroblasts exhibited high expression of RGS5, COX4I2 and FAM162B, whereas the MCAM− CAFs showed high expression of extracellular matrix components, such as FAP, PDGFRA and PDPN (Figure 4C). It has been reported that FAP+ CAFs are associated with an immunosuppressive microenvironment in multiple cancers(32). Additionally, PDPN expression in CAFs has been associated with increased risk of disease recurrence in lung adenocarcinoma(33). Therefore, our findings indicates that these MCAM− fibroblasts might contribute to an immunosuppressive microenvironment of SCLC.

Recently, a new ATOH1+ SCLC subtype was identified in an analysis of 38 SCLC CDX models(17). A novel PLCG2+ SCLC cell subpopulation with stem-like features was also identified and associated with exhausted CD8+ T cells and an immunosuppressive myeloid cell population(20). Interestingly, multiple hybrid SCLC cell subpopulations have been unmasked in this study. Hybrid or intermediate states of tumor cells have been widely reported in cancers, exemplified by the hybrid E/M cells(34). Moreover, hybrid Endothelial/Mesenchymal and hybrid Tumor/Endothelial cells have also been identified in cancers(35,36). Recently, hybrid tumor cells have been revealed by single-cell sequencing in several cancers(16,34). Phenotypic plasticity of SCLC has been proposed as a mechanism by which SCLC cells become more aggressive. For example, an intermediate state between SCLC-A and SCLC-N has been revealed in human SCLC(20). Moreover, a continuum of plastic cell states between SCLC subtypes was revealed by archetype analysis of bulk and scRNA-seq data(30). In addition, combined SCLC and NSCLC has been identified in a small number of lung tumors(29). For example, transformation from NSCLC to SCLC has become one of the mechanisms driving resistance to immune checkpoint inhibitors(37). In line with these observations, we also dissected the phenotypic plasticity of SCLC cells, exemplified by the presence of multiple hybrid SCLC cells, referring to tumor cells with multiple cell lineage features, within the heterogenous SCLC ecosystems.

For instance, we uncovered several novel hybrid cells including hybrid SCLC/LUAD and hybrid tumor/immune cells. Recently, tumor cells expressing immune signatures have been identified in several cancers such as nasopharyngeal carcinoma(21), skin squamous cell carcinoma(22), and head and neck squamous cell carcinoma(38). For example, a tumor cell subpopulation expressing dual markers of tumor and immune cells (EPCAM, MHC-II and complement genes) was revealed in nasopharyngeal carcinoma and observed to have increased tumorigenic properties and correlate with a poor prognosis(21). Moreover, fusion of tumor cells with leukocytes has been reported to contribute to tumor heterogeneity and increased metastatic behavior(39,40). However, to our knowledge, SCLC cells showing immune features have not been described in their ecosystems. We observed that several tumor cell subpopulations co-express markers of immune cells (CD45 and CD27) and neuroendocrine cells (INSM1 and ASCL1). We therefore postulate that these hybrid tumor/immune cells are more likely tumor cells with immune cell features. However, the biological roles of these hybrid cells will need to be determined in future studies.

Our study provides several novel findings. To start with, this is the first single-cell transcriptome landscape of cellular components and their spatial distributions in human primary SCLC ecosystems. Secondly, this study provides a comprehensive view and molecular signatures of distinct infiltrating immune cells, stromal fibroblasts and endothelial cells in human SCLC ecosystems. Third, we identify multiple hybrid states of SCLC cells, spanning from hybrid SCLC subtypes, hybrid SCLC/LUAD to hybrid tumor/immune cells, suggesting that dynamic evolution of SCLC cells occurs in SCLC ecosystems. Fourth, the evolution of these hybrid SCLC cells has also been revealed as well as new potential cell of origin for the SCLC-P subtype. Finally, this study offers a comprehensive view of the cell-cell communications within and across cellular components in human SCLC ecosystems, providing rationale to target these interactions for SCLC therapy. Given the heterogenous nature of SCLC, future studies will need to integrate more SCLC samples and incorporate higher resolution spatial transcriptomics technologies. It will also be critical to determine the functional roles and clinical relevance of the various hybrid states of tumor cell states that we have identified. In conclusion, this study provides a comprehensive single-cell atlas of cellular components and their spatial distributions in human SCLC ecosystems, revealing extensive heterogeneities of tumor, immune and stromal cells within tissues and tumors as well as insight into their transcriptional states. Our study also unmasked multiple hybrid tumor cell states in the human SCLC, indicating that targeting these hybrid tumor cells could form the basis for the next generation of therapeutic strategies for SCLC patients.

## METHODS

### Human tumor specimens

The human primary SCLC tumor and adjacent lung tissues were obtained from patients underwent surgery treatments in Shanghai Pulmonary Hospital. All patients provided signed informed consent. This study was conducted following the protocols approved by the Institutional Review Board at Shanghai Pulmonary Hospital (K21-344) and in accordance with the Declaration of Helsinki ethical guidelines. The pathology of all tumors was diagnosed by senior lung pathologists. The information of clinical specimens was provided in Supplementary Table 1. The representative histological images of tumors were provided in Supplementary Figure 7. Molecular subtypes of these tumors were defined by IHC staining of transcription factors ASCL1, NEURDO1, POU2F3 and YAP1 (Supplementary Figure 8). Notably, we observed that most tumors contain a mixed population of two or three SCLC subtypes with different proportions, including SCLC-A/SCLC-N, SCLC-A/SCLC-P and SCLC-P/SCLC-Y, suggesting that most primary SCLC tumors composed of a mixture of SCLC subtypes.

### Tumor dissociation

Fresh tumor specimens were transported to laboratory on ice and processed for the scRNA-seq and spatial transcriptomics analyses within 2 hours after surgical resection. Tumor tissues were washed 2 times with cold PBS and cut into small pieces (1-2 mm^3^). Next, tumor tissues were dissociated in 1 mg/mL Collagenase IV in PBS (Sigma-Aldrich) for 45 minutes at 37°C incubator. The digested tissue suspension was filtered sequentially by 70 μM and 40 μM strainers (Corning). Red blood cells (RBC) were removed from cell suspension using RBC lysis buffer (BioLegend). The dissociated cells were washed with PBS and resuspended in 1% FBS in PBS. The cell viability and concentration were determined by automated Cell Counter System (Invitrogen), only cell suspensions with viability above 85% were qualified for further scRNA-seq experiments.

### Single-cell RNA sequencing

The scRNA-seq libraries were constructed using 10X Genomics Chromium Single Cell 3′ v3 reagents (v3 Chemistry) or Chromium Next GEM Single cell 3′ v3.1 reagents (v3.1 Next GEM Chemistry) following the protocols provided by the manufacturer. In particular, we loaded around 12,000 cells to generate the library for each sample. The quality of libraries was analyzed by 2100 Bioanalyzer system (Agilent). All the libraries were sequenced on illumina Hiseq platform using PE150 sequencing mode. Sequencing depth was around 408 million reads/sample (∼30,412 reads/cell). The detailed information of sequencing was provided in Supplementary Table 2.

### scRNA-seq data processing

The Cell Ranger (version 4.0.0) was used to process reference genome alignment, quantify cells and genes with default parameters. The raw sequencing reads were mapped to human genome assembly GRCh38. The Seurat (3.2.0) R package was further used to process the gene-barcode counts matrices derived from the Cell Ranger output. Data from each of the 7 samples were merged into one Seurat object. The merged object was normalized and scaled by calculating total number of UMI count and Scaling Factor ratio of 10000 for individual cell. The following parameters were used to identify high-quality cells, including min.cells = 1, nFeature_RNA > 200 and mitochondrial gene fraction < 10%. The Scrublet package was used to predict and remove doublets(41). The batch effects were removed using the canonical correlation analysis (CCA) subspace alignment implemented in the Seurat. We also analyzed the individual sample separately in case that the batch correction may filter out some critical cells.

The principal component analysis (PCA) dimensionality reduction was run to visualize the similarity among cells using default parameters. Based on the PCA results, the top 30 principal components (PCs) were taken to perform graph-based clustering analysis, and the identified cell clusters were plotted by t-Distributed Stochastic Neighbor Embedding (t-SNE) and Uniform Manifold Approximation and Projection (UMAP) dimensionality reduction methods, respectively. According to the clustering analysis results, the overall correlation and PCA dimensionality reduction between cell clusters were determined. Subsequently, the *FindAllMarkers* function and *bimod* likelihood ratio test were employed to identify differentially expressed genes (DEGs) in different clusters (logfc.threshold = 0.1, min.pct = 0.01). Regarding the DEGs, genes that meet P value <= 0.05 and >= 1.5 times of the differential expression range were designated as marker genes for one cell cluster against the rest of cell clusters. For the detected marker genes, we further filtered out genes encoding ribosomal proteins, while kept genes that meet the following conditions: avg_logFC > 0.25, pct.1 > 0.25, pct.1 > pct. 2, p_val < 0.05. Finally, the top 10 genes of each cell cluster were displayed by heatmap, featureplot, vlnplot and dotplot, respectively. Cell types were annotated using canonical lineage markers for epithelial and lung cancer cells (EPCAM, NAPSA, SFTPB, SFTPC, SCGB1A1, INSM1, CGHA, SYP, ASCL1, NEUROD1, POU2F3, YAP1), immune cells (CD45, CD3D, CD4, CD8A, CD19, CD79A, CD20, CD138, CD14, CD68, HLA-DRA), endothelial cells (PECAM1, CD34, CDH5), and fibroblasts (ACTA2, COL1A1, FAP, PDGFRA, MCAM, PDPN).

### Spatial Transcriptomics analysis

The 10X Genomics Visium Spatial Transcriptomics platform was used to analyze the transcriptome of cellular components within the tissue context in 8 human primary SCLC tumors. The experiments were performed according to the protocol provided by the manufacturer. Briefly, fresh tumor tissues were embedded in OCT, snap-frozen in liquid nitrogen and stored at –80°C freezer. 10 μM thickness of tissue sections were prepared for subsequent experiments, including H&E staining, permeabilization and library construction. Specifically, all the tissue sections were permeabilized for 12 minutes. The barcoded libraries were sequenced using the illumina Nova Seq 6000 platform with the PE150 sequencing mode.

### Spatial transcriptomics data processing

The Space Ranger pipeline was used to process the Visium Spatial Gene Expression data following the guidelines provided by the manufacturer, including demultiplex, hg38 human reference genome alignment, tissue detection, fiducial detection, and barcode/UMI counting. The Visium spatial barcodes were further used to generate feature-spot matrices, determine clusters and perform gene expression analysis. Accordingly, high-quality spots (>= 200 genes/spot) were selected for subsequent analyses, including clustering analysis, marker gene analysis and functional annotation. In order to correct the sequence depth of high-quality spots, the *SCTransform* in Seurat was used to normalize and identify highly differential expression genes. The PCA was used to perform dimensionality reduction and compare the similarity among spots. The top 30 PCs of each spot were used for graph-based clustering analysis, and the identified clusters were plotted by t-SNE and UMAP, respectively. Based on the results of spot clustering by Seurat, the *FindAllMarkers* function in Seurat was used to identify DEGs for different spot clusters (logfc.threshold = 0.1, min.pct = 0.01). Accordingly, each spot cluster has its own marker genes, and genes that meet P value <= 0.05 and fold change >= 1.5 are designated as the marker genes between one cluster against all the remaining clusters. For the detected marker genes, we performed further filtering and selection: (1) filtered out genes encoding ribosomal proteins, (2) kept genes that meet the following conditions: avg_logFC > 0.25, pct.1 > 0.25, pct.1 > pct. 2, p_val < 0.05. Finally, the top 10 genes of each cluster were plotted by heatmap, featureplot, vlnplot and dotplot.

### Gene Ontology and KEGG enrichment analyses

For the DEGs identified in each cell cluster, the TopGO package was used to perform Gene Ontology (GO) enrichment analysis, whereas the KEGGREST package was employed to perform KEGG enrichment analysis. The Fisher’s exact test was used to calculate P value, and the P value was further examined by Benjamini & Hochberg’s test and the FDR was obtained.

### Trajectory analysis

Pseudotime trajectory analysis was performed by Monocle (2.12.0) as described previously(42). Based on the marker genes of each cluster, the DDRTree (0.1.5) was used to perform dimensionality reduction and construct a minimum spanning tree (MST). Next, the optimal ranking of single cells in high-dimensional and low-dimensional spaces was searched to generate the best cell differentiation trajectory curve. If the number of cells is more than 20,000, a representative 20,000 cells were randomly selected for the analysis. For instance, a representative 20,000 tumor cells were randomly selected for the trajectory analysis of all tumor cells extracted from the scRNA-seq data in this study. Besides, we first re-clustered all the cells of each cell type, then we performed trajectory analysis of the tumor cells, immune cells, fibroblasts and endothelial cells independently.

### Cell-cell interaction analysis

An in-house algorithm Cell Interaction Analysis (CellInter) was developed to infer potential ligand-receptor interactions between cellular components, including the cancer cells, T cells, B cells, Myeloid cells, fibroblasts and endothelial cells. Briefly, list of human ligand and receptor pairs was downloaded in the CellPhoneDB database (https://www.cellphonedb.org/downloads), which includes I2D, IMEx, InnateDB, InnateDB-All, IntAct, MINT, MatrixDB, MINT, and UniProt databases. The Seurat normalized expression matrix was used to perform the CellInter analysis. First, the product value of expression levels of paired ligand and receptor between two cell types was calculated, respectively. Next, the marker genes of each cell type were used to calculate the P value and FDR value using the Sample function in R with default 1000 times of permutations, respectively. FDR <= 0.05 was used to filter the ligand and receptor pair between two cell types. Accordingly, the number of ligand and receptor pair was counted between any two cell types and the diagram of interaction network among cell types was constructed. Specifically, the lines with direction in the diagram represent the number of interaction pairs between two cell types, whereby the arrows start from ligands and end at receptors. The color of each line is consistent with the cell type expressing the ligands. Moreover, the diagram of ligand and receptor interaction pairs among cell types was also generated, which displayed the top pairs of interaction based on the rank of calculated score (average expression of ligand and receptor) of each pair.

### Copy-number variation analysis

The InferCNV (https://github.com/broadinstitute/infercnv) was used to infer copy number alterations of single cells from the scRNA-seq data as described previously(43). Default parameters were used. In particular, T cells were used as reference to infer copy number variations in other cell types. A heatmap diagram was generated to display the relative gene expression intensities across individual chromosome.

### Histological and immunohistochemical staining

The human SCLC tumor and adjacent lung tissues, xenograft tumors were fixed in NBF and embedded in paraffin. 4 μM thickness sections were prepared for hematoxylin and eosin (H&E) or immunohistochemical (IHC) staining using standard protocols. The primary antibodies used for the IHC staining included: anti-ASCL1 (1:100, Rabbit polyclonal, Abcam), anti-NEUROD1 (1:100, Rabbit polyclonal, Abcam), anti-YAP1 (1:100, Rabbit polyclonal, Cell Signaling Technology), and anti-POU2F3 (1:250, Rabbit polyclonal, NB1-83966, Novus). The representative photos of H&E and IHC staining were taken using Leica DM6 B microscope.

### Cell lines

Human SCLC cell lines (NCI-H1048, NCI-H1694, NCI-H1092 and NCI-H211) were purchased from ATCC. NCI-H1048, NCI-H1694 and NCI-H1092 were cultured in DMEM/F12 media supplemented with 10% FBS (Gibco) and 1% Pen/Strep (Gibco). NCI-H211 was cultured in RPMI-1640 media supplemented with 10% FBS and 1% Pen/Strep. All cell lines were cultured in cultured at 37 °C in a 5% CO2 incubator and regularly checked for mycoplasma contamination.

### Flow cytometry

The flow cytometry was performed using standard protocols. Human SCLC cells were labeled with the following antibodies: BV421 Mouse anti-human CD45 (BD, 563879), PE mouse anti-human CD8 clone RPA-T8 (BD, 561949), and APC mouse anti-human CD27 clone M-T274 (BD, 561400). BV421 Mouse IgG1 k Isotype Control (BD, 562438), PE Mouse IgG1 κ Isotype Control (BD, 555749) and APC Mouse IgG1 κ Isotype Control (BD, 555751) antibodies were used as isotype control. The experiment was repeated twice independently.

### Statistical analyses

All statistical analyses and graphical representation of data were performed in the R environment or using GraphPad Prism 9.0 software.

## Supporting information

Supplementary Figures 1-6, related to Figure 1-7

Supplementary Figures 7-8, related to Human tumor specimens in Methods section

Supplementary Tables 1-4, related to Figure 1, 2 and 7

## Authors’ Contributions

DJ conceived and designed this project, wrote the manuscript. DJ and YZ supervised this project. JC and YZ collected the tumor specimens and clinical information. DJ, HO, CH, LS, WJ and DP performed experiments. DJ, CX and CH performed data analysis. JL analyzed the pathology of tumors. AA edited the manuscript.

## Acknowledgements

This work was supported by NSFC (82072571), Shanghai Pujiang Scholar Program (19PJ1408500), Shanghai Young Eastern Scholar Program, Experimental Animal Research Fund, Science and Technology Commission of Shanghai Municipality (19140905600), Shanghai Jiao Tong University School of Medicine Scientific and Technological Innovation Program (19XJ11009) and Shanghai Jiao Tong University School of Medicine Full-time Clinical Research Team Program. This work was partly supported by the dream mentor-outstanding Young Talent Program (fkyq1903) and Shanghai Pulmonary Hospital Program (fkzr2114). We thank Dr. David MacPherson (Fred Hutch) for critical reading of the manuscript. We thank Genergy Bio-Technology (Shanghai) Co. for providing sequencing service and helping with data analyses.

## Supplementary Figure Legends

**Supplementary Figure 1**

A, Copy number variations of each cell cluster were analyzed based on scRNA-seq data. T cell clusters 11 and 12 were used as reference.

B, UMAP visualization of all cell clusters in each of the 7 SCLC samples, respectively.

C, UMAP visualization of all cells colored by chemotherapy treated and untreated samples.

D, UMAP visualization of all cell clusters in chemotherapy treated and untreated samples, respectively.

**Supplementary Figure 2**

A, Images of H&E staining and spatial spot clusters for 6 SCLC tissue regions, respectively.

B, Spatial feature plots of clusters and their distribution in 7 SCLCs, respectively.

C, Inter- and intra-tumoral heterogeneity revealed by Spatial Transcriptomics in 8 SCLCs.

D, Heatmap showing top 10 upregulated genes in each spatial spot cluster of LC-251 tissue.

E, Spatial feature plots showing the hybrid SCLC/LUAD cells within LC-51 tissue.

**Supplementary Figure 3**

A, The relative proportions of 23 immune cell clusters in each of the 7 SCLCs was illustrated.

B, UMAP visualization of 23 immune cell clusters in each of the 7 SCLCs, respectively.

C, KEGG pathway enrichment analysis of CD20+ B cells in SCLC ecosystems.

D, Spatial feature plots showing the expression of representative markers for B cells and myeloid cells within the LC-251, LC-250 and LC-29 tumor tissues, respectively.

**Supplementary Figure 4**

A-B, Top enriched KEGG pathways identified in S1 and S2 fibroblasts, respectively.

C, The expression levels of ASCL1, GRP and INSM1 in each state of fibroblasts.

D-E, Violin plots showing expression of representative genes in each cluster of fibroblasts.

F, The expression levels of ASCL1, GRP, INSM1 and HLA-DRA in each state of ECs.

G, Dot plot showing the expression of ANGPT2, ESM1 and RGS5 in each state of ECs.

H, Spatial feature plots of representative stromal cell markers within LC-251 and LC-250 tissues, respectively.

**Supplementary Figure 5**

A, UMAP visualization of 26 tumor cell clusters in each of the 7 SCLCs, respectively.

B, UMAP plots showing the expression of representative markers in all tumor cells.

C, Flow cytometry results showing the proportion of tumor cells expressing each of the immune cell markers (CD45, CD27 and CD8) in human NCI-H1882 and NCI-H1092 cell lines, respectively.

D, UMAP plot showing the expression of endothelial cell markers in tumor cells.

E-F, Spatial feature plots showing the localization of hybrid SCLC subtypes within the LC-4 and LC-29 tissues, respectively.

**Supplementary Figure 6**

A, Expression of KRT5, KRT8, NKX2-1 and SFTPB in each state projected onto pseudotime space in LC-29 tissue.

B, Dot plot showing the top interactions between endothelial and tumor cells. Dot colors refer to the total mean of individual partner average expression values, dot sizes represent the - log10(p values + 1e-04).

C, UMAP plots showing the expression of CD74, MIF, NRP1, VEGFB, TIMP1 and FGFR2 in all cellular components in 7 SCLCs.

**Supplementary Figure 7**

Representative images of H&E staining for 11 human primary SCLC tumors. Scale bar, 100μM.

**Supplementary Figure 8**

Representative images of IHC staining for ASCL1, NEUROD1, POU2F3 and YAP1 protein levels in 11 human primary SCLC tumors. Scale bar, 100μM.

## Supplementary Tables

Supplementary Table 1 Clinical information of 12 human SCLC specimens.

Supplementary Table 2 Summarization of scRNA-seq and spatial transcriptomics sequencing information.

Supplementary Table 3 List of ligand-receptor interactions across 27 cell clusters in 7 SCLC ecosystems.

Supplementary Table 4 List of ligand-receptor interactions across 8 spatial spot clusters within the LC-251 tissue.

