## Supplementary Figures 1-6, related to Figure 1-7 for "Single-Cell Transcriptomic Analyses of Tumor Ecosystems and Spatial Architectures in Human Small Cell Lung Cancer"

### Supplementary Figure 1

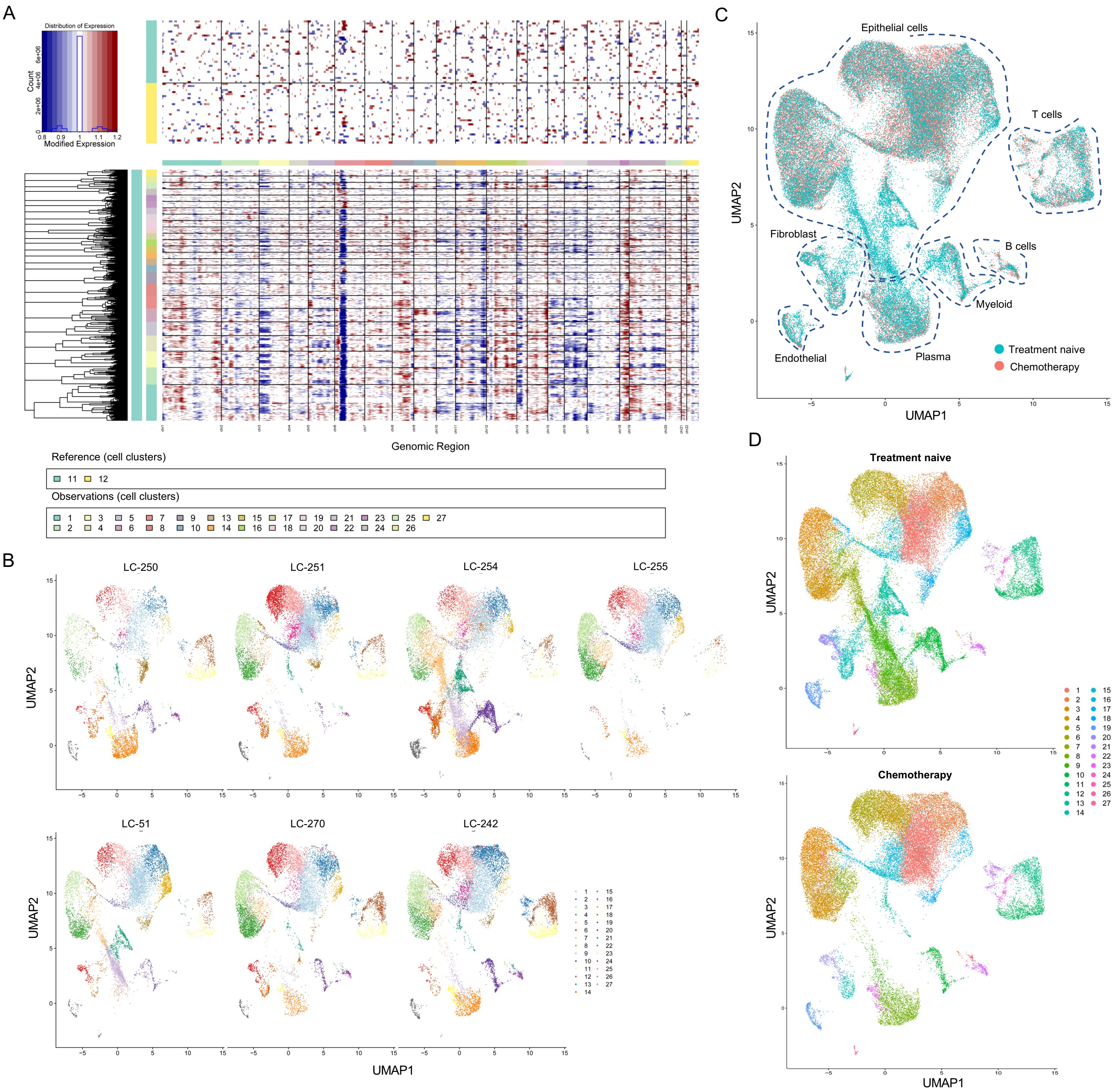

### Supplementary Figure 2

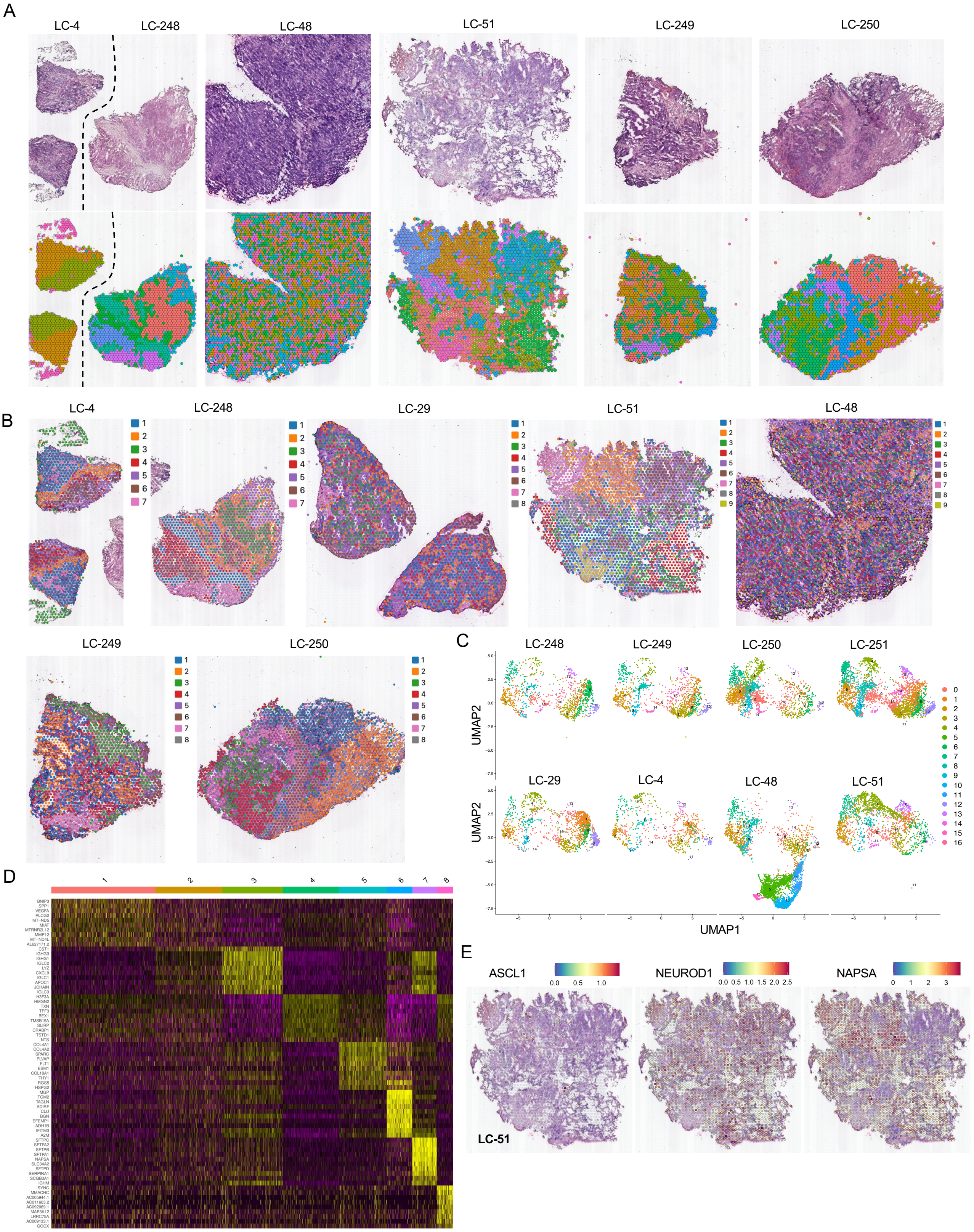

### Supplementary Figure 3

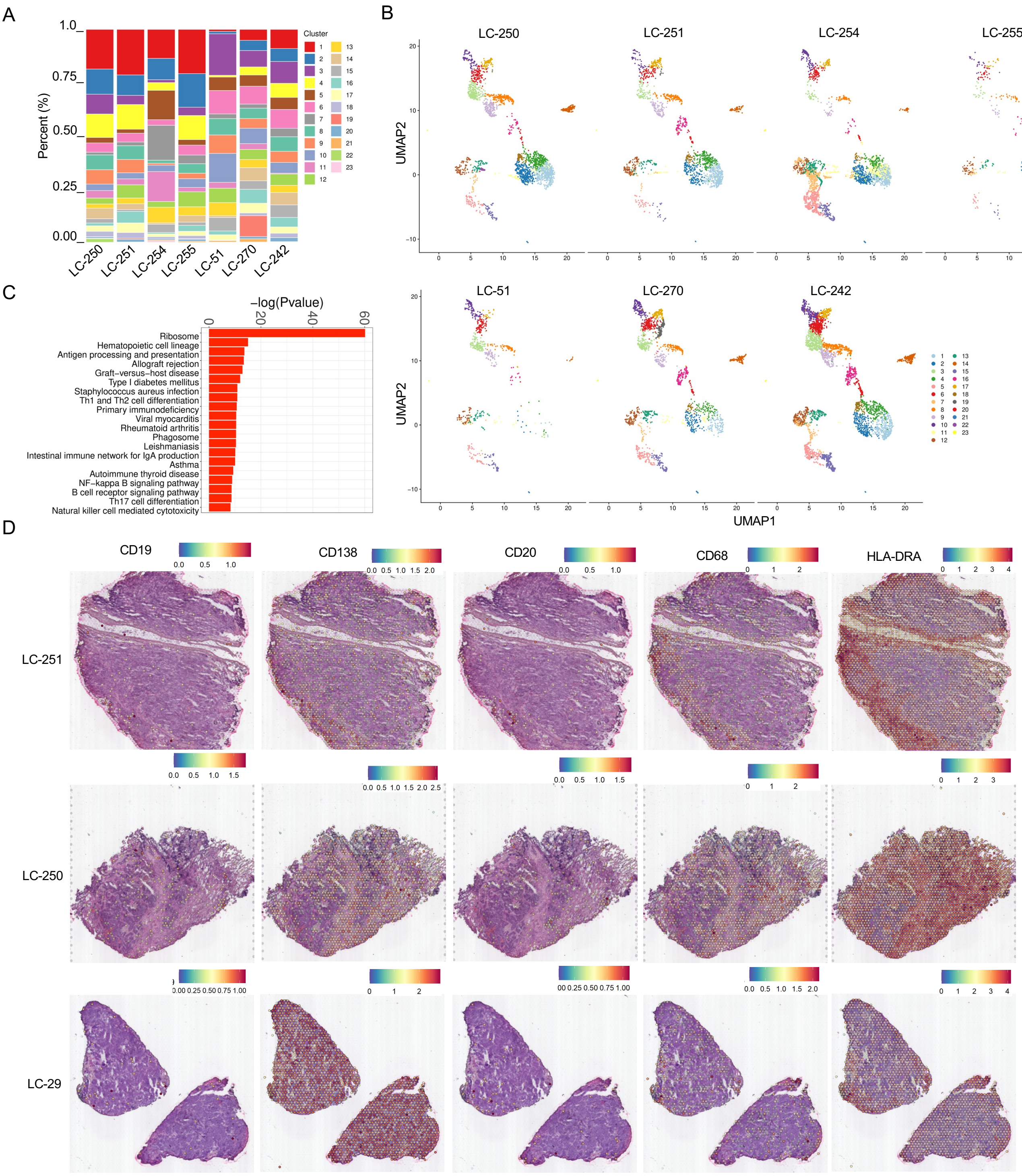

### Supplementary Figure 4

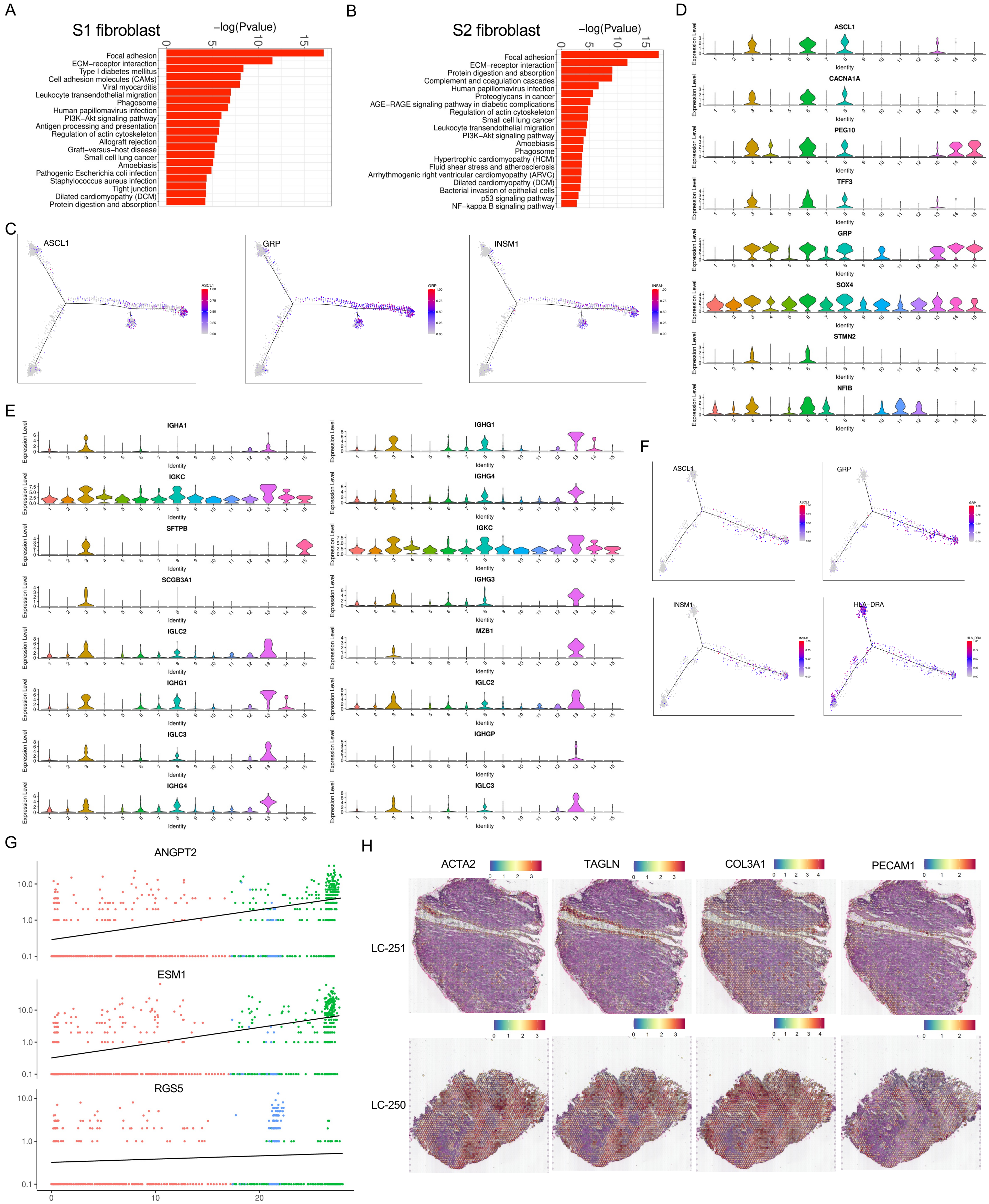

### Supplementary Figure 5

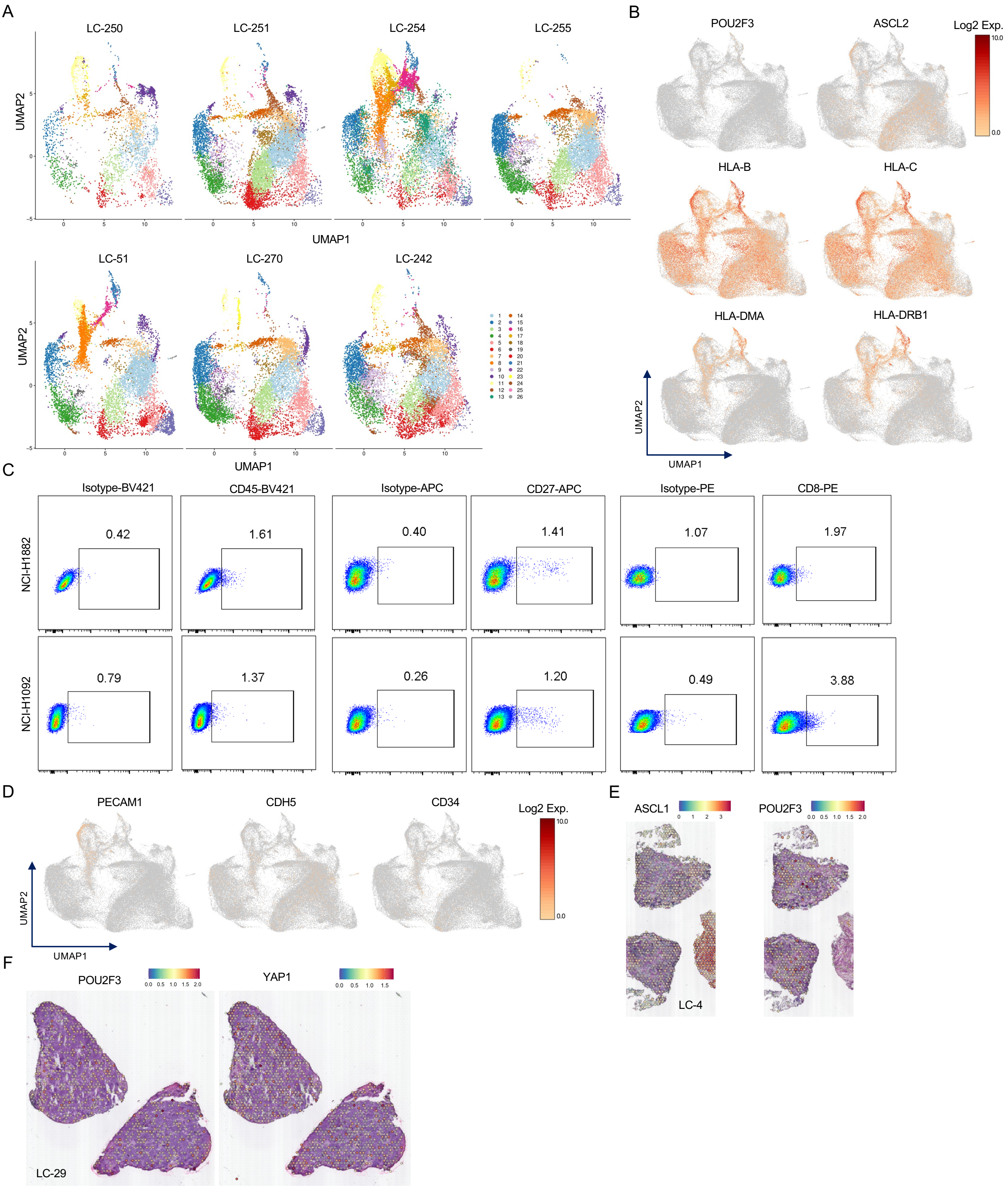

### Supplementary Figure 6

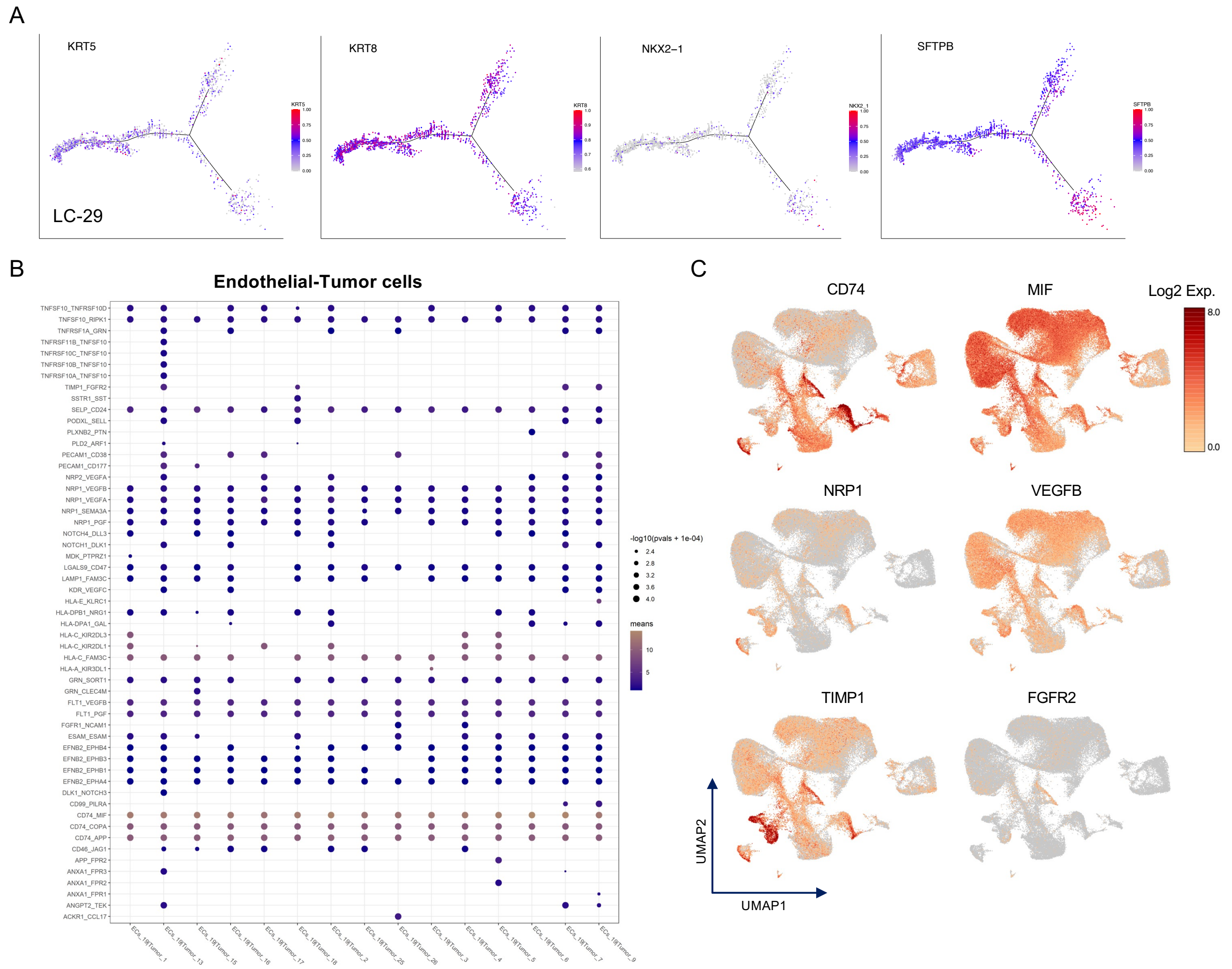
