## Supplementary figures and images for "Single-Cell Transcriptomic Analyses of Tumor Ecosystems and Spatial Architectures in Human Small Cell Lung Cancer"

### Supplementary Figures 7-8, related to Human tumor specimens in Methods section

#### Supplementary Figure 7

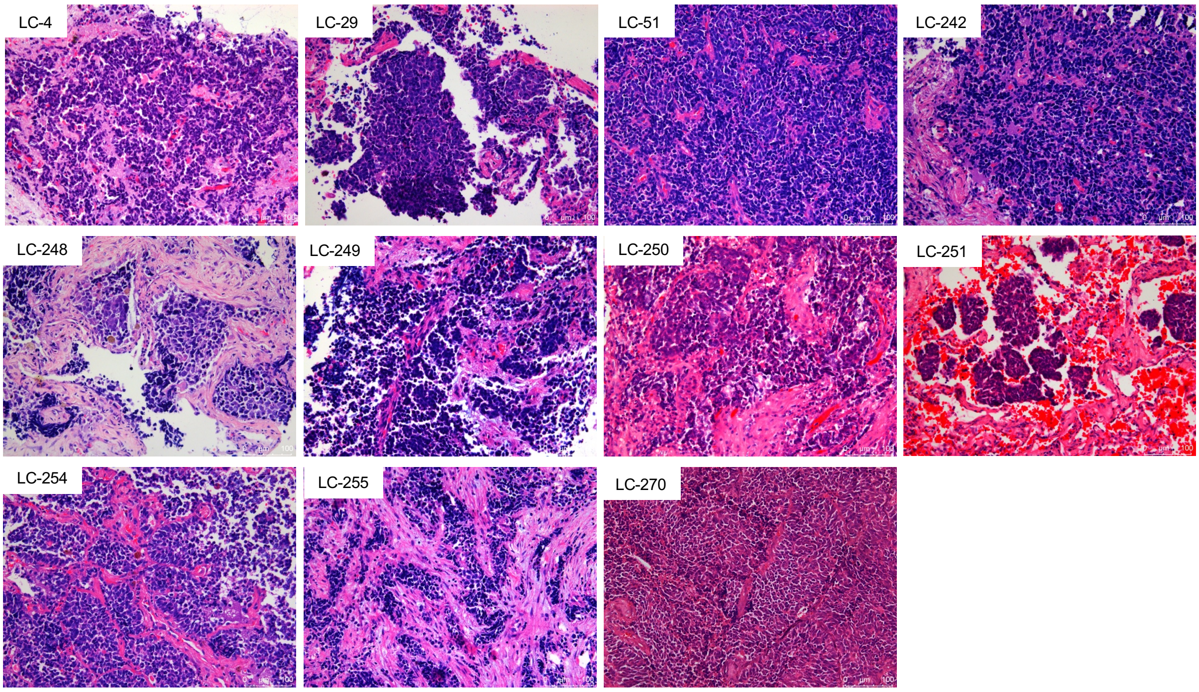

Supplementary Figure 8

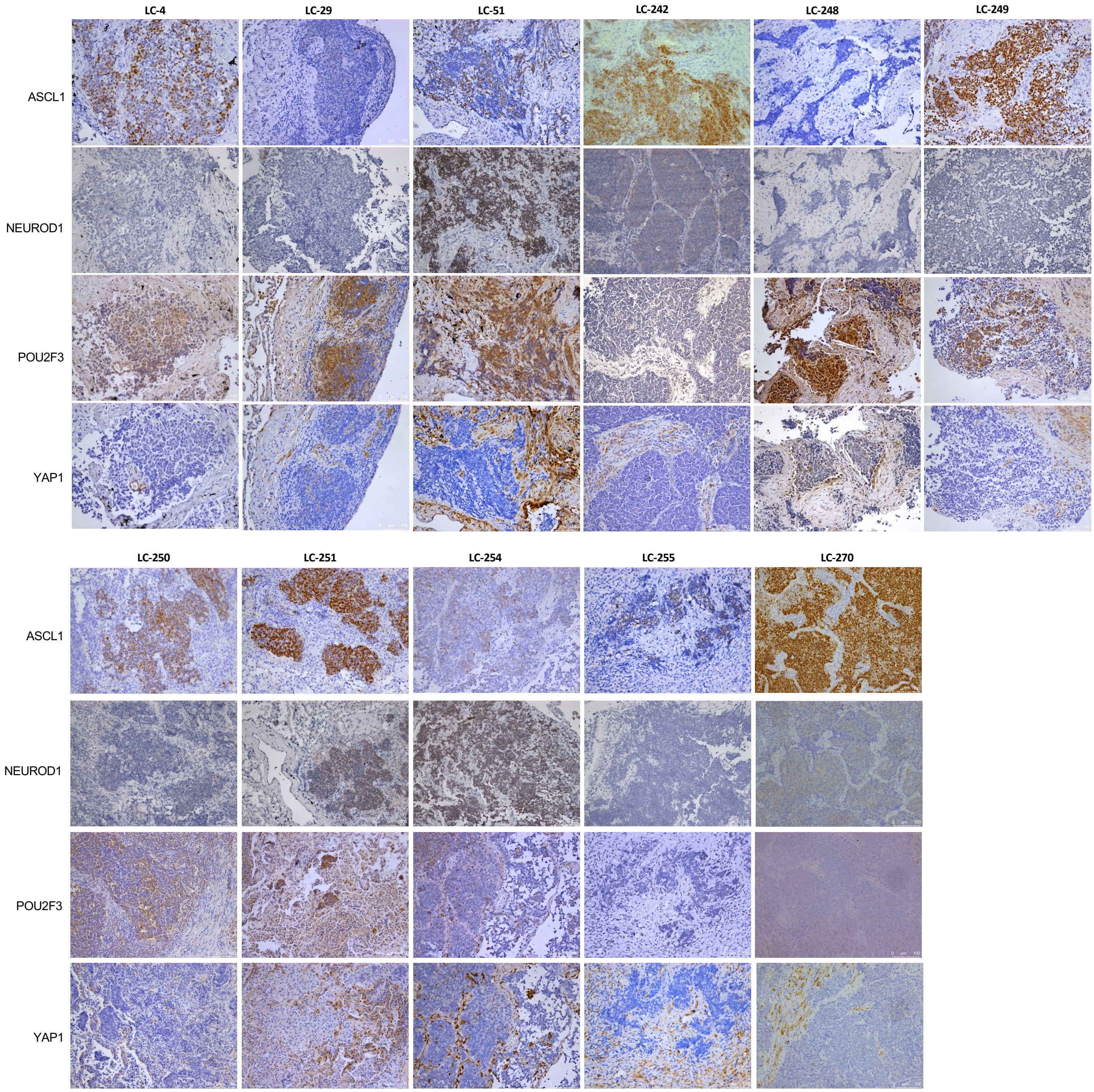
